# Pan-cancer analysis of single-cell RNA sequencing data from 304 human tumors sheds light on the ‘aneuploidy paradox’

**DOI:** 10.64898/2026.05.06.723237

**Authors:** Guy Wolf-Dankovich, Tomer Mashiah, Ron Saad, Einav Somech, Haia Khoury, Itay Tirosh, Uri Ben-David

**Author notes:** Corresponding author: Uri Ben-David. **Conflict of interest:** U.B.-D. receives consulting fees from Accent Therapeutics and research funding from Galmed Pharmaceuticals. R.S. is a current employee of CytoReason. I.T. is a scientific advisor of Immunitas therapeutics and Compugen, and is a co-founder and advisor of Cellyrix therapeutics.

## Abstract

Aneuploidy poses a central paradox in cancer biology: it impairs cellular fitness in normal cells but drives cancer progression. To resolve this, we analyzed single cell transcriptomes from >665,000 cells – including ∼288,000 malignant cells – across 304 tumors and 15 cancer types. Integrating transcriptomics with inferred aneuploidy profiles, we characterized cell-intrinsic programs and interactions with the tumor microenvironment. Unexpectedly, highly aneuploid single cells exhibited reduced proliferation and metabolism, contrasting sharply with tumor-bulk profiles. We show this divergence is driven by karyotypic heterogeneity: in highly heterogeneous tumors, aneuploid cells display signatures of acute stress and negative selection. Conversely, in clonally aneuploid tumors, these detrimental signatures are lost and replaced by signatures of increased proliferation and enhanced metabolism, reflecting adaptation. Additionally, we identified consistent transcriptional programs driven by recurrent chromosome-arm alterations across both single cells and bulk tumors. These findings illuminate the selective forces shaping tumor evolution and the aneuploidy paradox.

**Statement Of Significance:** By jointly evaluating gene expression and aneuploidy at single-cell resolution, we demonstrate that karyotypic heterogeneity underlies the transcriptional impact of aneuploidy, reveal distinct cellular responses to emerging versus stable states, and identify recurrent chromosomal alterations that drive conserved transcriptional programs. Our findings capture the dynamics of aneuploidy evolution in human tumors, providing novel insights into the ‘aneuploidy paradox’.

## Introduction

Aneuploidy, the presence of an abnormal number of chromosomes, is one of the most prevalent alterations in human cancer (1), yet its functional role remains paradoxical (2). On one hand, tumors with high levels of aneuploidy are frequently associated with poor prognosis, increased proliferation, and immune evasion (3,4). On the other hand, studies in model organisms and in normal and premalignant human cells have shown that aneuploidy imposes a fitness cost, impairing proliferation and triggering stress responses (5–7). This apparent contradiction – commonly referred to as the aneuploidy paradox (2) – suggests that the impact of aneuploidy is highly context-dependent, and is likely shaped by the tumor environment, evolutionary pressures, and the timing of karyotypic change (1).

Resolving this paradox requires moving beyond static measurements of aneuploidy, characterizing the dynamics and heterogeneity of chromosomal alterations within individual human tumors (8–10). Single-cell RNA sequencing (scRNA-seq) offers a powerful framework to address this challenge by enabling joint inference of gene expression and chromosomal copy number at the cellular level (11,12). Unlike tumor-bulk analyses that average signals across populations, single-cell data capture the coexistence of distinct karyotypic states within tumors, revealing both newly emerging and clonally selected aneuploidies (13). This can allow us to study how individual cells respond to aneuploidy, how selection shapes the transcriptome of aneuploid clones, and whether karyotypic diversity modulates tumor behavior.

Recent large-scale single-cell analyses suggest that genetic subclones – defined by copy number profiles – may explain only a limited fraction of the transcriptional heterogeneity observed within tumors (14–16). These findings point to a potentially important role for non-genetic factors, including cellular plasticity, epigenetic regulation, and microenvironmental context, in shaping tumor cell states. Nevertheless, the recurrent nature of specific chromosome-arm alterations across diverse tumor types (3,11,17), strongly suggests that these events are subject to positive selection, very likely due to the gene expression changes that they induce. Therefore, identifying consistent transcriptional programs that are activated by recurrent clonal and subclonal aneuploidies is key to understanding the biological impact of recurrent aneuploidies and the selective forces that shape tumor karyotypes.

In this study, we analyzed ∼288,000 malignant cells, mapping inferred aneuploidy profiles onto single-cell gene expression data to yield a final cohort of approximately 247,000 cells with matched aneuploidy and gene expression profiles across 304 tumors spanning multiple cancer types. We examine how aneuploidy levels and karyotypic heterogeneity shape gene expression at both the single-cell and tumor-bulk levels, revealing distinct transcriptional patterns associated with emerging versus clonally stable karyotypes. We then systematically investigate the impact of recurrent chromosome-arm alterations, identifying consistent gene expression changes across single cells and tumors that point to their role in driving tumor evolution and selection. Finally, we leverage this high-resolution data to characterize the surrounding tumor microenvironment, investigating how aneuploidy correlates with the composition and abundance of specific immune cell types and revealing an enrichment of M2 tumor-associated macrophages (TAMs) in aneuploid tumors.

## Results

### Generating matched single-cell gene expression and chromosome-arm copy number calls for 304 human tumors

To analyze the impact of aneuploidy levels and heterogeneity, we used scRNA-seq data from >665K single cells, including ∼288K malignant cells, spanning 304 tumors across 36 studies (Fig. 1A). These data were obtained from the Curated Cancer Cell Atlas (14,18). Using copy-number calls inferred from the scRNA-seq data with inferCNA (12), we determined the aneuploidy status of each chromosome-arm of every individual cell (Methods; Supplementary Table S1), resulting in matched gene expression and aneuploidy profiles for ∼247K malignant single cells.

**Figure 1:**
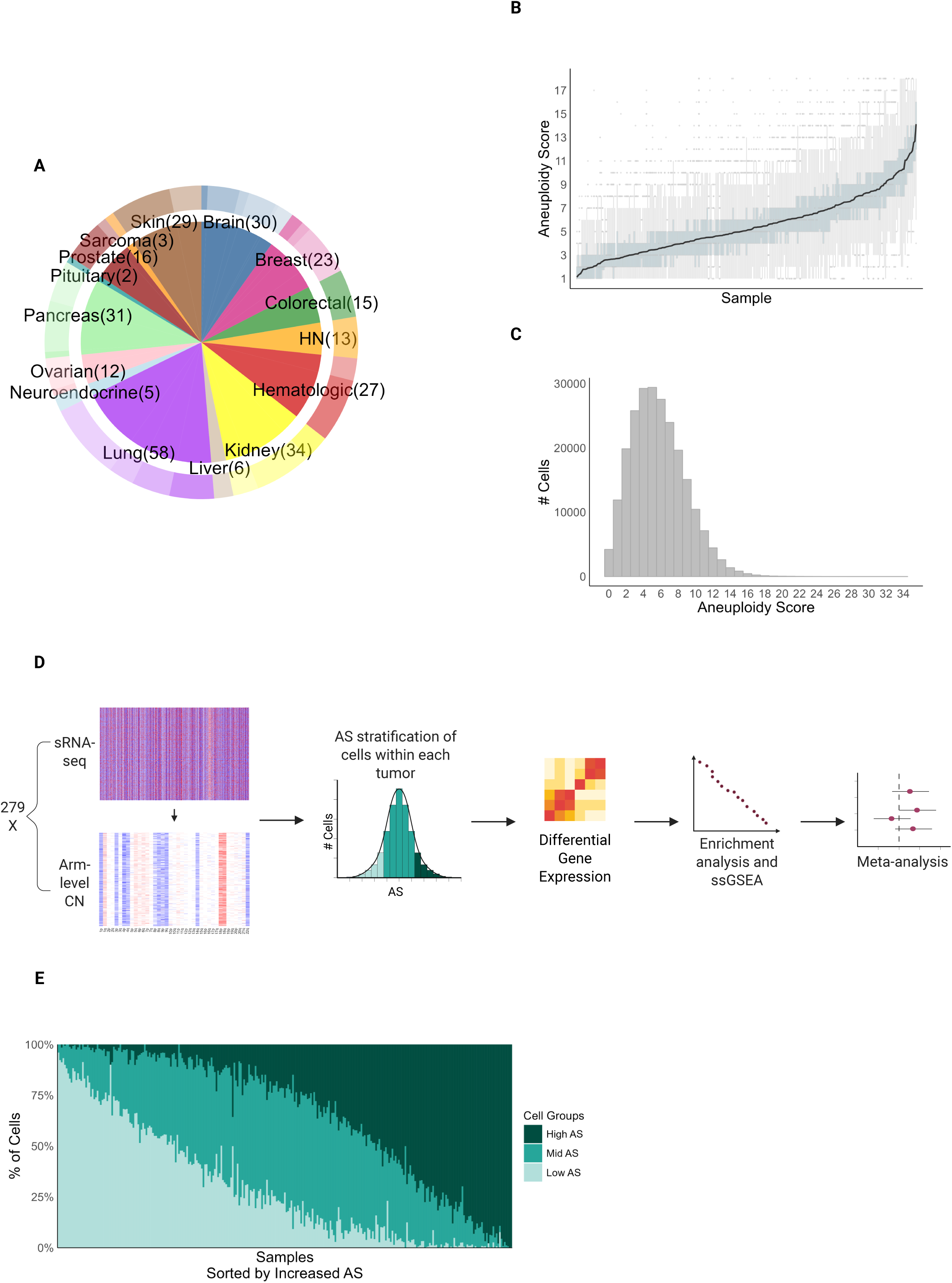
Overview of datasets and analysis workflow. **A)** Distribution of Cancer Types in the Dataset: Pie chart summarizing the representation of 15 distinct cancer types in the compiled dataset. The sizes of the slices are proportional to the number of published studies that contributed tumor samples for each cancer type (n = 304). Colors represent different cancer types, and the number of source publications for each type is indicated in parentheses. **B)** Distribution of Aneuploidy Scores (AS) Across Tumor Samples: Bar plot displaying the AS values for each of the 279 tumor samples, ordered by increasing AS along the X-axis. The Y-axis represents the tumor-level AS. Background box plots illustrate the distribution of malignant single-cell AS values within each corresponding tumor. **C)** Distribution of Malignant Cells by Aneuploidy Score: Histogram representing the number of malignant cells (total n = 234,744 across 279 tumor samples) as a function of their individual AS. The X-axis shows the AS, while the Y-axis indicates the frequency of cells at each score across the entire dataset. **D)** Illustration of the analytical workflow. Key steps include generating a matched gene expression and arm-level copy number dataset, stratifying the malignant cells by aneuploidy score, comparing high-AS vs. low-AS cells within the same tumor using differential gene expression analyses and linear regressions, and performing gene set enrichment analyses and meta-analyses across the tumor cohort. Created with BioRender.com. **E)** Stratification of cells into AS groups: Bar plot illustrating the relative distribution of cells across three AS-defined categories: Low (AS ≤ 3, in light green), Mid (3 < AS < 7, in medium green), and High (AS ≥ 7, in dark green). Each bar corresponds to a tumor (n = 279), with the proportion of cells belonging to each AS group color-coded accordingly. The tumor samples are ordered by increasing AS.

As a first step of quality control, we examined the number of cells and the total number of genes expressed within each tumor (Supplementary Fig. S1A-B). Of 304 tumors, 279 had sufficient numbers of cells and expressed genes (>2,000 expressed genes and >100 single cells with copy number data), and were used for our downstream analyses. Using the chromosome-arm level aneuploidy calls, we defined the Aneuploidy Score (AS) for each cell as its number of aneuploid chromosome arms (19), and determined the overall AS for each tumor by averaging these scores across all of its malignant cells. Our dataset comprises tumors with varying levels of aneuploidy (Fig. 1B,C), ranging from highly aneuploid tumors to those with near-diploid karyotypes (see examples in Supplementary Fig. S1C-F). Additionally, we observed a broad range of aneuploidy levels within tumors, as many tumors contained both cells with near-diploid karyotypes and those with high(er) aneuploidy levels (see examples in Supplementary Fig. S2).

Using this rich and diverse dataset, we set out to examine the transcriptional effects of aneuploidy levels and of karyotypic heterogeneity, at the cell-level and at the tumor-level, with and without clonal expansion. Recognizing the previously reported associations between aneuploidy and immune infiltration (3,4), we also sought to explore with our data the effect of aneuploidy on tumor cell composition.

### Aneuploidy is associated with opposite gene expression patterns in malignant single cells and in bulk tumors

We first set out to study the cell-intrinsic effects of aneuploidy within the malignant compartment. To investigate the transcriptional consequences of aneuploidy, we implemented a comprehensive analysis pipeline for each of the 279 tumor samples (Method; Fig. 1D).

We first used the matched gene expression and aneuploidy profiles of each malignant cell to categorize the individual cells into three groups based on aneuploidy scores (AS): Low (AS ≤ 3), Mid (3 < AS < 7), and High (AS ≥ 7). Among our 279 tumors, many (∼40%) included malignant cells from both Low and High groups, while the rest primarily consisted of cells from just one of these groups (Fig. 1E). For downstream single-cell-based analyses, we focused on the 114 tumors that had ≥30 malignant cells from both the Low-AS and High-AS groups. These tumors represent a variety of cancer types (13 of the 15 cancer types included in the dataset; Fig. 1E and Supplementary Fig. S1G).

Next, to compare High-AS vs. Low-AS cells within each tumor, we conducted Differential Gene Expression (DGE) and Pathway Enrichment (PE) analyses in three different ways, in order to explore the intra-tumor interactions between aneuploidy levels and gene expression programs. We then examined recurrent patterns across tumors in order to evaluate whether AS has a consistent impact on transcriptional alterations. Surprisingly, our comparison of High-AS to Low-AS cells revealed that high aneuploidy is often associated with decreased proliferation and metabolic activity, as well as an elevated inflammatory response (Fig. 2A, Supplementary Fig. S3A-B and Supplementary Table S2). These findings appear to contrast with earlier tumor-balk observations, which indicated that tumors with high levels of aneuploidy tend to be more aggressive, proliferative, and exhibit less immune activity (1,3,4).

**Figure 2:**
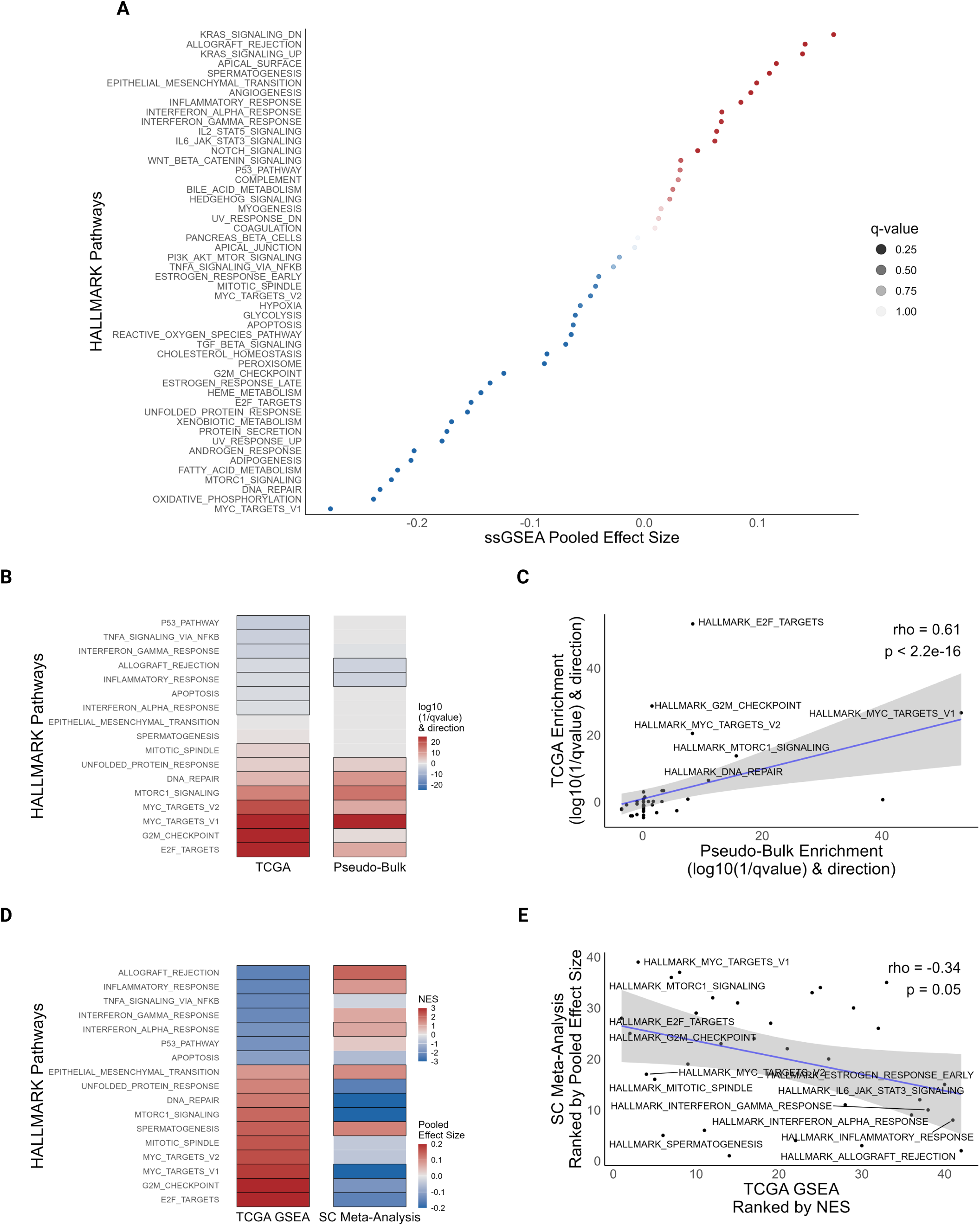
Aneuploidy is associated with opposite gene expression patterns in single cells and in bulk tumors. **A)** ‘Hallmark’ gene set enrichment analysis in single cells: Single-sample GSEA (ssGSEA) meta-analysis comparing High-AS vs. Low-AS cells across multiple tumors (n=114). The X-axis shows pooled effect sizes; the Y-axis lists ‘Hallmark’ pathways. Dot color indicates direction of enrichment (red = upregulated; blue = downregulated in High-AS), and transparency reflects statistical significance (q-value). **B)** ‘Hallmark’ gene set enrichment analysis in bulk tumors: Enrichment analysis comparing High-AS vs. Low-AS tumors in both pseudo-bulk (aggregated single-cell) and TCGA datasets. Shown are selected ‘Hallmark’ pathways related to proliferation, metabolism, and immunity. Heatmap color represents both the direction and strength of enrichment, with red indicating upregulation and blue indicating downregulation in High-AS tumors. Black boxes mark significantly enriched pathways (q < 0.05). **C)** Concordant pathway enrichment between bulk datasets: Scatter plot comparing enrichment direction and strength between pseudo-bulk and TCGA tumors. The x-axis represents pseudo-bulk; the y-axis represents TCGA. A significant positive correlation is observed (Spearman’s rho = 0.61, p < 2.2e–16), indicating overall agreement between the analyses. **D)** Contrasting pathway patterns in single-cell vs. bulk tumors: Selected ‘Hallmark’ gene sets related to proliferation, metabolism, and immunity are shown across TCGA (left) and single-cell (right) datasets, comparing High-AS vs. Low-AS tumors or cells, respectively. Color represents enrichment results (NES for TCGA; Polled effect size for single-cell). Significant pathways (q < 0.05) are highlighted with a bold border. **E)** Inverse correlation between single-cell and bulk tumor pathway changes: Scatter plot comparing ranked pathway enrichments between single-cell and TCGA datasets. The x-axis ranks TCGA ‘Hallmark’ pathways (Normalized Enrichment Scores); the y-axis ranks the same pathways by single-cell pooled effect sizes. A significant negative correlation is observed (Spearman’s rho = –0.34, p = 0.05), suggesting opposite expression trends at the single-cell and bulk levels.

To determine whether these differing findings are unique to our dataset composition, we conducted a pseudo-bulk gene expression data analysis of the 279 tumors, comparing High-AS tumors to Low-AS tumors. We then compared these results with a bulk-level gene expression data analysis of The Cancer Genome Atlas (TCGA) cohort. Interestingly, High-AS tumors in our dataset showed increased expression of proliferation and metabolic programs, along with reduced inflammatory response, in contrast to our single-cell analysis (Fig. 2B, Supplementary Fig. S3C and Supplementary Table S3), and the same trend was also observed in the TCGA cohort (as was previously reported (3)). Additionally, we found a strong positive correlation between our pseudo-bulk level analysis and the TCGA results (Fig. 2C). This correlation held true and even became more significant when accounting for tumor purity in both cohorts (Supplementary Fig. S3D-E and Supplementary Table S3).

Our results therefore reveal an intriguing inverse relationship between pathway changes observed at the single-cell vs. the tumor-bulk levels (Fig. 2D and Supplementary Fig. S3F), with opposite trends of pathway activation and suppression (up-regulation or down-regulation) in the two types of analysis (Fig. 2E).

### Transcriptional signatures of aneuploidy are robust to technical artifacts and cell cycle confounding

Given these unexpected findings, we implemented an additional layer of quality control to evaluate whether the observed trends might be driven by technical artifacts. We quantified the number of expressed genes and the mitochondrial transcript fraction for each cell as proxies for cellular fitness and sequencing quality (20). Initially, we hypothesized that cells of the Low-AS group might be of lower quality, as low cell quality could potentially diminish the power of aneuploidy detection. Contrary to this concern, we found that the High-AS cells actually possessed a significantly lower mean number of expressed genes and a higher mitochondrial transcript fraction compared to the Low- and Mid-AS populations (Supplementary Fig. S4A-B).

As a reduction in these metrics can often signal low-quality libraries or apoptotic cells, we sought to determine if this trend represented a genuine biological consequence of aneuploidy. To address this, we implemented a stringent filtering pipeline, excluding cells with fewer than 1,500 expressed genes or a mitochondrial fraction exceeding 10%. Notably, even after the removal of these lower-quality outliers, we observed a highly significant reduction in the number of expressed genes with increased aneuploidy (Supplementary Fig. S4C). Furthermore, the equivalent levels of mitochondrial reads following filtering (Supplementary Fig. S4D) suggests that there is no inherent technical quality difference in cell sequencing between the groups. Instead, these findings are consistent with the reduced proliferation and metabolism observed in the aneuploid cells. Finally, upon re-conducting the single-cell DGE and PE analyses on this high-quality cell subset, the previously observed trends in High-AS versus Low-AS cells remained consistent (Supplementary Fig. S4E-F), confirming the robustness of these findings.

To ensure that our findings were independent of the specific parameters employed for aneuploidy inference, we performed a sensitivity analysis using more stringent scoring criteria. This alternative approach was specifically designed to mitigate the detection of “false positive” events, which are inherent to single-cell analyses. Indeed, some aneuploidy events were (presumably falsely) detected within the non-malignant cell populations (Supplementary Fig. S2). We therefore repeated the aneuploidy calling with more stringent calling thresholds (Methods), effectively minimizing these false-positive events across the dataset (Supplementary Fig. S5). While this conservative approach carries the inherent risk of excluding genuine aneuploidies in certain malignant populations, it provides a stringent baseline for our functional comparisons.

We then repeated the gene expression analyses with cells classified based on these stringent parameters (Supplementary Fig. S6A-C). Importantly, the results were highly similar to those observed with the more lenient aneuploidy calling parameters: High-AS cells continued to exhibit decreased proliferation and metabolic activity alongside an elevated inflammatory response (Supplementary Fig. S6D), while the pseudo-bulk analysis maintained the inverse trend of increased proliferative and metabolic signaling (Supplementary Fig. S6E-H). These results demonstrate that the divergence between the single-cell and tumor-bulk transcriptional patterns associated with aneuploidy is a robust biological phenomenon that persists whether the aneuploidy calling algorithm is tuned to minimize false positives or false negatives.

As a final quality control step, we addressed the potential confounding effect of the cell cycle. In highly proliferative populations, cell-cycle genes can dominate the transcriptome and represent a major source of non-genetic variation in scRNA-seq (21). Because copy number inference relies on averaged expression signals, it is sensitive to such coordinated transcriptional programs, which can bias the inferred genomic profiles (22). To determine whether differences in cell cycle phase could explain the observed gene expression changes, we performed the pseudo-bulk and single-cell-based analyses separately for non-cycling cells (G1 phase) and cycling cells (S/G2/M phases; Methods). To ensure robust inference, we restricted this analysis to tumors retaining a minimum of 100 malignant cells from the relevant phases. In both comparisons, we still observed a contrasting relationship between High-AS and Low-AS cells across multiple proliferative and metabolic pathways (Supplementary Fig. S7), indicating that the cell cycle is not a major contributor to the observed contradiction.

Taken together, these extensive validation and quality control measures confirm that the observed transcriptomic shifts are inherent to the aneuploid state and are not driven by technical artifacts, thresholding biases, or cell cycle distributions.

### Karyotypic heterogeneity is associated with reduced proliferation, p53 activation, and negative selection

To shed light on the inverse relationship between the aneuploidy-associated pathway changes in the single-cell level and in the tumor-bulk level, we examined several potential explanations.

The first explanation for this contradiction relates to tumor cell composition: while bulk-level analyses incorporate all tumor cells, our single-cell analysis focused exclusively on malignant cells. To investigate whether this distinction was responsible for the observed differences, we conducted a pseudo-bulk level analysis using only the malignant cells. The contrasting relationship was retained for multiple pathway changes (Supplementary Fig. S8A), indicating that this is not the culprit of the apparent contradiction.

Next, we considered that the tumor cohort composition might contribute to the observed contradiction: while the bulk-level analyses utilize all tumors from our dataset (n=279) or from the TCGA cohort, including all aneuploidy groups, our single-cell analysis specifically focused on tumors that contained both High-AS and Low-AS cells (n=114; Supplementary Fig. S8B). Consequently, the single-cell analysis included more tumors with mid-level aneuploidy (Mid-AS). To determine whether this difference in dataset composition contributed to the observed differences in gene expression, we conducted a pseudo-bulk level analysis using only the tumors included in the single-cell analysis. As most of the tumors included in this analysis belonged to the Mid-AS group, the differences between the more aneuploid tumors and the less aneuploid tumors were weaker than in the original analysis, and many pathways were not altered at all. Nonetheless, this did not revert the gene expression signatures in the tumor-bulk analysis, and a couple of proliferative and metabolic pathways (MYC and the unfolded protein response) remained upregulated in the aneuploid tumors although strongly downregulated in the highly aneuploid single cells (Supplementary Fig. S8C).

Having ruled out these potential confounders, we turned to explore biological explanations. We hypothesized that the difference between the single-cell and tumor-bulk analyses might reflect the differences in the effect of emerging aneuploidy vs. adapted aneuploidy. The single-cell analysis allows us to analyze ‘snap shots’ that include both emerging karyotypes and those that have already been selected for. When analyzing tumors with both High-AS and Low-AS cells, we could distinguish between two types of tumors: (1) those that display stable aneuploid clones with low overall heterogeneity; and (2) those with heterogeneous karyotypes. Therefore, we hypothesized that the highly-aneuploid cells that arise within karyotypically heterogeneous tumors would show evidence of negative selection (reflected by lower proliferation and metabolism and elevated inflammatory response), whereas those that are observed in more clonal tumors have already been through positive selection (reflected by higher proliferation and metabolism and reduced inflammatory response). This could explain the apparent contradiction between the single-cell analysis, which considers (by definition) mostly heterogeneous tumors, and the bulk-level analysis, which considers mostly High-AS tumors with clonal aneuploidies.

To put this hypothesis to test, we next explored the intra-tumor karyotypic heterogeneity in our dataset. Using Pearson’s correlation, we calculated pairwise similarities between the chromosome-arm aneuploidy state of the malignant cells. We averaged the cell-to-cell correlations within each tumor to come up with karyotypic heterogeneity scores. A high mean correlation indicates that the tumor has low karyotypic heterogeneity. Conversely, a low mean correlation signifies dissimilarity of aneuploidy patterns, implying high karyotypic heterogeneity. As expected, tumors containing both High-AS and Low-AS cells demonstrated a greater degree of karyotypic heterogeneity (Supplementary Fig. S8D), in line with our hypothesis.

Next, we explored the relationship between aneuploidy levels and karyotypic heterogeneity. Interestingly, we found a strong negative correlation: tumors with lower aneuploidy levels exhibited greater heterogeneity, while those with higher aneuploidy levels were more karyotypically homogeneous (Fig. 3A and Supplementary Fig. S8E-F). The negative correlation between aneuploidy levels and karyotypic heterogeneity was also evident when the analysis was limited to the tumors containing both High-AS and Low-AS cells (Fig. 3B).

**Figure 3:**
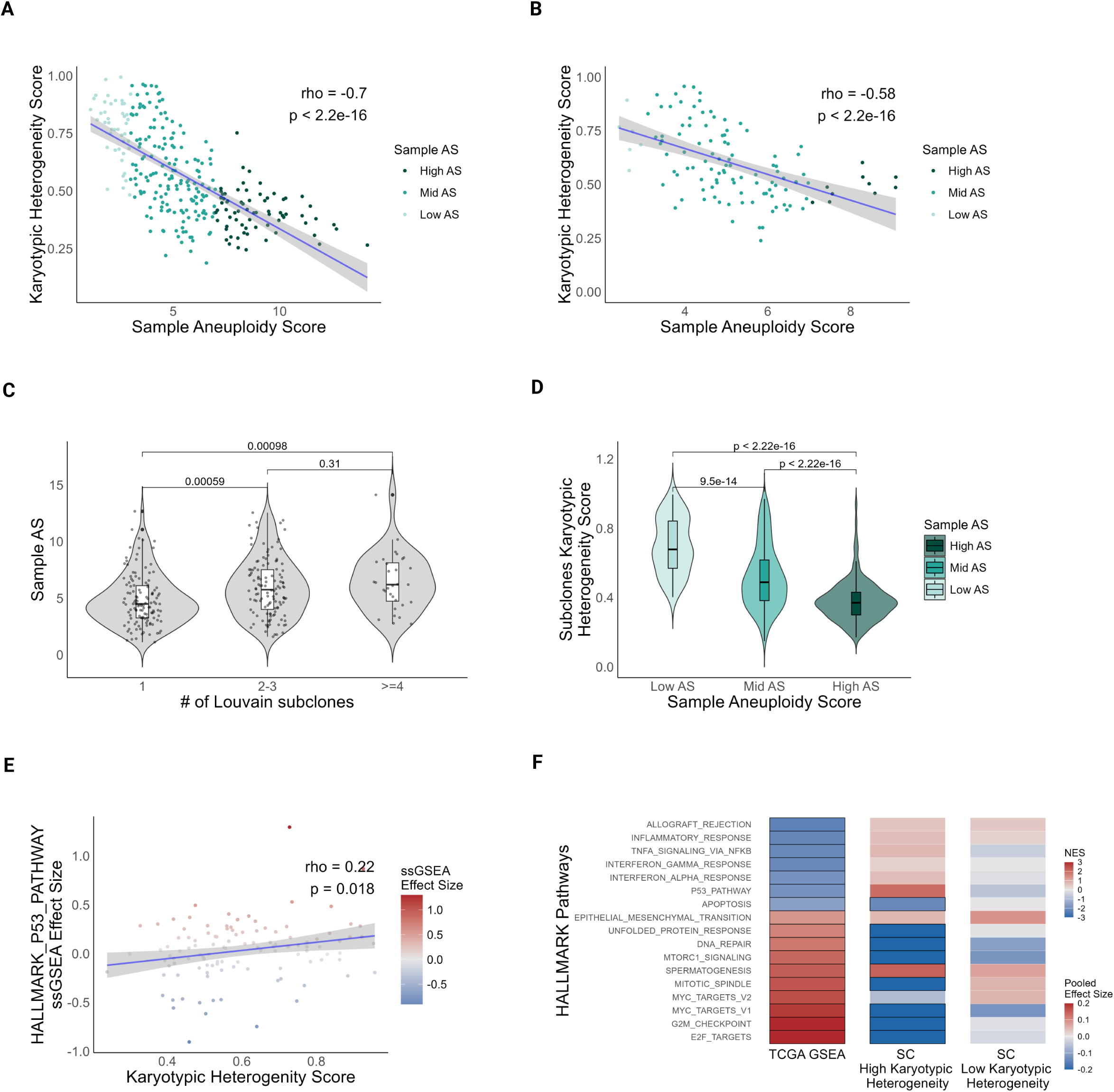
Aneuploidy heterogeneity is associated with reduced proliferation, p53 activation and negative selection. **A)** Negative correlation between aneuploidy scores and karyotypic heterogeneity scores: Scatter plot showing the relationship between tumor aneuploidy score (AS, x-axis) and karyotypic heterogeneity score (y-axis). Each point represents a tumor (n=279), color-coded by AS group: Low (light green), Mid (medium green), and High (dark green). A strong negative correlation is observed (Spearman’s rho = –0.7, p < 2.2e–16). **B)** Negative correlation between AS and karyotypic heterogeneity scores in tumors comprising both High-AS and Low-AS cells: Same as (A), but analysis restricted to tumors with cells from both the High-AS and the Low-AS groups (n=114). A significant negative correlation remains (Spearman’s rho = –0.58, p < 2.2e–16). **C)** Aneuploidy levels across genetic subclones: Violin plots showing tumor AS stratified by the number of genetic subclones (1, 2–3, or ≥4), as defined by Louvain clustering of copy number profiles. Tumors with a single subclone exhibit significantly lower AS compared to those with 2–3 (p = 0.00059) or ≥4 subclones (p = 0.00098; two-tailed Wilcoxon rank-sum test). No significant difference was observed between the 2-3 and ≥4 subclone groups (p = 0.31). **D)** Aneuploidy heterogeneity across genetic subclones: Violin plots showing karyotypic heterogeneity scores of genetic subclones, stratified by tumor AS group. Subclones from High-AS tumors show significantly lower heterogeneity compared to Mid-AS (p < 2.22e–16) and Low-AS groups (p < 2.22e–16). Additionally, Mid-AS subclones exhibit significantly lower heterogeneity than Low-AS subclones (p = 9.5e-14; two-tailed Wilcoxon rank-sum test). **E)** Correlation between p53 activity and karyotypic heterogeneity: Scatter plot showing the relationship between tumor-level karyotypic heterogeneity (x-axis) and the ssGSEA effect size of the ‘Hallmark’ p53 gen set (y-axis), comparing High-AS vs. Low-AS cells. Each dot represents a tumor (n=114), dot colors indicate AS group: Low (blue), Mid (gray), and High (red). A significant positive correlation is observed (Spearman’s rho = 0.22, p = 0.018), suggesting that higher heterogeneity is associated with increased p53 activation. **F)** Pathway enrichment in heterogeneous vs. homogeneous tumors: Selected ‘Hallmark’ gene sets related to proliferation, metabolism, and immunity are shown for High-AS vs. Low-AS comparisons in both TCGA (left) and single-cell (SC) data. In SC data, results are stratified by tumor karyotypic heterogeneity, comparing the most heterogeneous one-third of tumors (middle, n=43) and the most homogeneous one-third (right, n=24). Colors represent enrichment results (NES for TCGA; pooled effect size for SC). Significant pathways (q < 0.05) are highlighted with a bold border.

To better understand this negative correlation, we investigated the association of aneuploidy levels with the number of genetic (copy number-defined) subclones. Using the Louvain Clustering approach (14,15), we defined genetic subclones and quantified the number of subclones per tumor (Supplementary Fig. S9A-E; Methods). We found that tumors with higher aneuploidy levels tended to comprise a greater number of genetic subclones (Fig. 3C). This observation supports previous reports linking increased aneuploidy to greater intratumoral heterogeneity (8,23,24). We then calculated the karyotypic heterogeneity for each subclone and examined its relationship with the overall aneuploidy levels of the corresponding tumor. Interestingly, we found that although High-AS tumors tend to have more subclones, these subclones are significantly more karyotypically homogeneous. In contrast, Low-AS tumors tend to have fewer subclones, with greater karyotypic heterogeneity (Fig. 3D). Taken together, our findings indicate that Low-AS tumors contain on average fewer subclones, but the aneuploid cells that are observed in these tumors are less similar to each other, representing emerging aneuploidies that are likely to get selected against. Conversely, High-AS tumors contain more subclones on average, and the aneuploid cells observed in these tumors are more similar to each other, reflecting stable karyotypes that have already been selected for.

Finally, when examining the activation of the p53 pathway, which protects cells from aneuploidy when functional (25–28), we observed a significant (albeit weak) positive correlation between the degree of karyotypic heterogeneity in tumors and the p53 pathway activity in the aneuploid cells within these tumors (Fig. 3E). In contrast, High-AS tumors exhibited lower p53 pathway activity than Low-AS tumors (Fig. 2B). This suggests that in karyotypically heterogeneous tumors, the p53 pathway is activated upon chromosome missegregation and aneuploidy formation, in line with their reduced proliferation, reduced metabolic activity and increased immune response; in contrast, in more homogeneous High-AS tumors, p53 is no longer activated, reflecting the positive selection of the clonal karyotypes as well as the well-established association between high degree of aneuploidy and p53 pathway silencing (3,29).

We conclude that the single-cell analysis focused on karyotypically heterogeneous tumors and therefore reflected the transcriptional consequences of emerging, temporary karyotypes. On the other hand, the tumor-bulk analyses focused on more karyotypically-homogeneous tumors and therefore reflected the transcriptional consequences of selected, stable karyotypes. In agreement with this conclusion, we observed an inverse relationship between the single-cell and tumor-bulk analyses of the most karyotypically heterogeneous tumors, but this inverse relationship was no longer observed in karyotypically homogeneous tumors (Fig. 3F, Supplementary Fig. S9B and Supplementary Table S2). These results confirm that karyotypic heterogeneity is indeed a major contributor to the gene expression analyses: in heterogeneous tumors, aneuploid cells are less proliferative and metabolically active than their diploid counterparts and exhibit a higher immune response; however, this is not the case in clonally aneuploid tumors, in which the aneuploid cells already represent adaptive karyotypes.

### Recurrent chromosome-arm alterations drive consistent expression changes across cells and tumors

Genetic clones within tumors are often defined by their karyotypes (30,31), yet it remains debatable to what extent these genetic clones are associated with unique gene expression patterns and phenotypes (14). We therefore examined whether genetic subclones within a tumor are systematically associated with activation/repression of specific signaling pathways. We used pathway enrichment analyses to compare each subclone to all other subclones of the same tumor, and assessed the number of pathways that were significantly enriched or depleted. Our findings revealed that, for both ‘Hallmark’ and ‘Reactome’ gene sets, the vast majority of subclones (∼80% and ∼93%, respectively) showed significant alterations in at least one pathway (a median of ∼7 and ∼93 pathways, respectively; Supplementary Fig. S10). These results indicate that karyotype-defined genetic subclones are characterized by distinct pathway-level transcriptional programs, but this does not necessarily mean that it is the specific karyotype that determines these transcriptional differences.

Next, we investigated the unique transcriptional changes associated with each specific, recurrent aneuploidy, in order to gain a better understanding of the positive selection for specific aneuploidies. We first identified recurrent chromosome-arm gains and losses in each cancer type based on the TCGA cohort (Methods; Supplementary Table S4) (17). We then compared the gene expression patterns between tumors and cells with and without each recurrent aneuploidy, utilizing the TCGA cohort and our collection of 279 tumors, respectively. Finally, we looked for consistent gene expression alterations to identify transcriptional changes linked to recurrent aneuploidies in each cancer type (Supplementary Tables S5-6).

In our investigation of recurrent chromosome-arm losses, we compared the expression of single cells with and without each recurrent chromosome-arm loss within the same tumor. Pathway enrichment analysis revealed that, on average, 17 ‘Hallmark’ pathways were differentially expressed between cells with vs. without recurrent chromosome-arm losses (Supplementary Fig. S11A). Furthermore, considering both ‘Reactome’ and ‘Hallmark’ pathways, we identified consistent dysregulation of specific signaling pathways between the single-cell and bulk cell-population level analyses for most chromosome-arm/cancer-type (CA-CT) pairs (Fig. 4A and Supplementary Fig. S11B).

**Figure 4:**
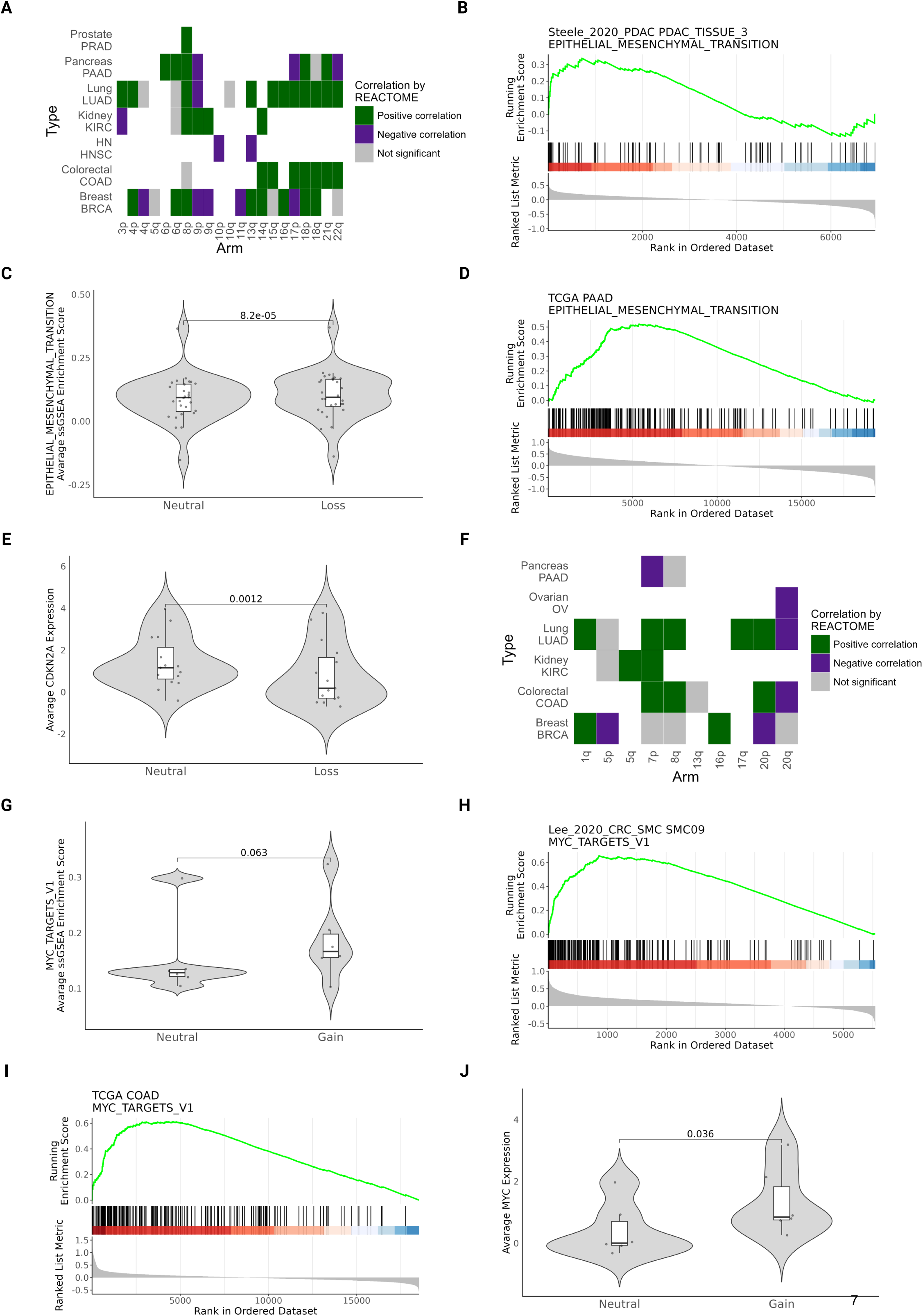
Recurrent chromosome-arm alterations drive consistent expression changes across cells and tumors. **A)** Concordant transcriptional changes induced by chromosome-arm losses across tumors and single cells: Heatmap showing the correlation between transcriptional changes in TCGA and single-cell datasets, assessed using ‘Reactome’ gene sets . Data spans 7 cancer types and 21 recurrent chromosome-arm losses (totaling 57 chromosome arm–cancer type pairs). Colors indicate correlation direction: green, significant positive correlation, purple, significant negative correlation, gray, not significant. **B)** Example GSEA plot showing EMT enrichment in a pancreatic tumor with Chr9p loss: Gene Set Enrichment Analysis (GSEA) plot for a representative pancreatic tumor, demonstrating EMT pathway upregulation in Chr9p-loss cells vs. Chr9p-WT cells. **C)** EMT pathway is upregulated in Chr9p-loss cells: Violin plots comparing ‘Hallmark’ Epithelial-Mesenchymal Transition (EMT) gene set enrichment scores (via ssGSEA) between Chr9p-loss and Chr9p-wild-type (WT) cells. Each point represents a within-tumor group average (n = 28 tumors; kidney and pancreas cancer types). The EMT gene set is significantly enriched in Chr9p-loss cells (paired two-tailed Wilcoxon signed-rank test, p = 8.2e–05). **D)** EMT pathway is upregulated in TCGA pancreatic tumors with Chr9p loss: GSEA plot of TCGA PAAD samples comparing the EMT gene set between Chr9p-loss vs. Chr9p-WT tumors. Enrichment is consistent with single-cell findings, indicating EMT upregulation in Chr9p-loss tumors. **E)** *CDKN2A* expression is reduced in Chr9p-loss cells: Violin plots comparing average *CDKN2A* expression between Chr9p-loss and Chr9p-WT cells across 15 tumors (kidney and pancreas). A significant reduction in *CDKN2A* expression is observed in Chr9p-loss cells (paired two-tailed Wilcoxon signed-rank test, p = 0.0012). **F)** Concordant transcriptional changes induced by chromosome-arm gains across tumors and single cells: Heatmap showing the correlation between transcriptional changes in TCGA and single-cell datasets, assessed using ‘Reactome’ gene sets. Data spans 6 cancer types and 10 recurrent chromosome-arm gains (totaling 25 chromosome arm–cancer type pairs). Colors indicate correlation direction: green, significant positive correlation, purple, significant negative correlation, gray, not significant. **G)** MYC pathway is upregulated in Chr8q-gain cells in colorectal tumors: Violin plots comparing ‘Hallmark’ MYC_targets_V1 gene set enrichment scores (via ssGSEA) between Chr8q-gain and Chr8q-WT cells. Each point represents a within-tumor group average (n = 6 colorectal tumors). The MYC gene set is enriched in Chr8q-gain cells (paired two-tailed Wilcoxon signed-rank test, p = 0.063). **H)** Example GSEA plot showing MYC pathway enrichment in a colorectal tumor with Chr8q gain: Gene Set Enrichment Analysis (GSEA) plot for a representative colorectal tumor, demonstrating MYC pathway upregulation in Chr8q-gain cells vs. Chr8q-WT cells. **I)** MYC pathway is upregulated in TCGA colorectal tumors with Chr8q gain: GSEA plot of TCGA COAD samples comparing the MYC gene set between Chr8q-gain vs. Chr8q-WT tumors. Enrichment is consistent with single-cell findings, indicating MYC upregulation in Chr8q-gain tumors. **J)** *MYC* expression is increased in Chr8q-gain cells: Violin plots comparing average *MYC* expression between Chr8q-gain and Chr8q-WT cells across 6 colorectal tumors. A significant increase in *MYC* expression is observed in Chr8q-gain cells (paired two-tailed Wilcoxon signed-rank test, p = 0.036).

One prominent example is the recurrent chromosome-arm loss of chromosome 9p (Chr9p). In both kidney and pancreatic cancer, we noted consistent dysregulations of ‘Hallmark’ pathways at the single-cell level, within tumors exhibiting mixed Chr9p copy number statuses, as well as at the tumor-bulk level, between tumors that differ in their Chr9p copy number status (Supplementary Fig. S11B). Specifically, we found an upregulation of the epithelial-mesenchymal transition (EMT) ‘Hallmark’ pathway in Chr9p-loss cells in both cancer types (Fig. 4B-C and Supplementary Fig. S11C-D). At the tumor-bulk level, we also observed upregulation of the EMT pathway for both cancer types (Fig. 4D and Supplementary Fig. S11E). Interestingly, previous studies have linked Chr9p loss to aggressiveness and metastasis (32), in line with the observed upregulation in EMT in tumors and cells that have lost this chromosome-arm. Furthermore, we noted a significant downregulation of *CDKN2A*, a candidate driver for Chr9p loss (17), in the Chr9p-loss cells (Fig. 4E and Supplementary Fig. S11F). Indeed, previous studies have shown that *CDKN2A* inhibits EMT, and its loss can disrupt the maintenance of epithelium, contributing to a more aggressive cancer phenotype (33,34).

Another interesting example is found at chromosome 18q (Chr18q) loss, which was associated with consistent gene expression dysregulation in colorectal, lung, and breast cancers at both tumor-bulk and single-cell levels (Fig. 4A). Investigating potential driver genes revealed that *SMAD2* and *SMAD4*, located on Chr18q, had been suggested as drivers in lung and colorectal cancers (17,35,36). In these cancer types, we observed significant downregulation of *SMAD2* and *SMAD4* at the single-cell level in Chr18q-loss cells within tumors exhibiting mixed Chr18q copy number statuses, as well as in the tumor-bulk level in tumors with this chromosome-arm loss (Supplementary Fig. S12). These genes are part of the transforming growth factor-β (TGFβ) signaling pathway (37,38), and our analysis at both the single-cell and tumor-bulk levels indeed indicated downregulation of several ‘Reactome’ pathways related to TGFβ signaling in the Chr18q-loss cells (Supplementary Fig. S13).

In our investigation of recurrent chromosome-arm gains we found that, on average, 19 ‘Hallmark’ pathways were differentially expressed between cells with vs. without the recurrent chromosome-arm gains (Supplementary Fig. S14A). As with the recurrent chromosome-arm losses, we identified consistent dysregulations of specific signaling pathways between single-cell and tumor-bulk results for most CA-CT pairs (Fig. 4F and Supplementary Fig. S14B).

A clear example is that of chromosome 8q (Chr8q) in colorectal cancer, where we observed an upregulation of MYC-related pathways in the Chr8q-gain cells within tumors exhibiting mixed Chr8q copy number statuses (Fig. 4G-H and Supplementary Fig. S14C) and in tumors with gain of that chromosome arm (Fig. 4I). The *MYC* oncogene is located on chromosome 8q, and Chr8q gain is known to be associated with its increased expression, which promotes tumorigenesis (39–41). Our findings indeed indicated a significant overexpression of *MYC* in cells with a single extra copy of Chr8q (Fig. 4J and Supplementary Fig. S14D), as well as in the Chr8q-gain TCGA COAD tumors (Supplementary Fig. S14E).

Similarly, we noted an upregulation of Notch-related signaling pathways in breast cancer cells with chromosome 1q (Chr1q) gain, within tumors with mixed Chr1q copy number statuses (Supplementary Fig. S14F-H) and in TCGA BRCA tumors with that chromosome-arm gain (Supplementary Fig. S14I-J). These findings are in line with several studies that have suggested that Chr1q gain could lead to an upregulation of Notch signaling in breast cancer (42,43).

In conclusion, our findings reveal that specific chromosome-arm gains and losses are associated with significant and recurring changes in gene expression, at both the single-cell and tumor-bulk levels. The same pathways that are dysregulated in single cells that harbor a specific aneuploidy within tumors that are heterogeneous for the status of that aneuploidy, are similarly dysregulated in tumors that clonally harbor that aneuploidy. These changes contribute to the distinct transcriptional programs that characterize different genetic subclones within the same tumor. This suggests that certain aneuploidies confer fitness advantages, influencing the characteristics of tumor cells and driving the selection of beneficial karyotypes.

### Single-cell resolution reveals immune microenvironment dynamics hidden in tumor-bulk analyses

Having established a framework for malignant-cell aneuploidy analysis, we investigated its association with the surrounding non-malignant compartment. Based on the 3CA cell annotations (14,18), we assigned cell type identity to single cells from 273 tumors (Methods; Fig. 5A and Supplementary Table S7). Across the cohort, T cells, macrophages, and fibroblasts emerged as the most abundant non-malignant cell types, in line with the previous reports (14). Because the microenvironmental architecture of hematological malignancies fundamentally differs from that of solid tissues, we excluded these samples from subsequent analyses. Our downstream investigation was therefore restricted to the remaining 248 solid tumors.

**Figure 5:**
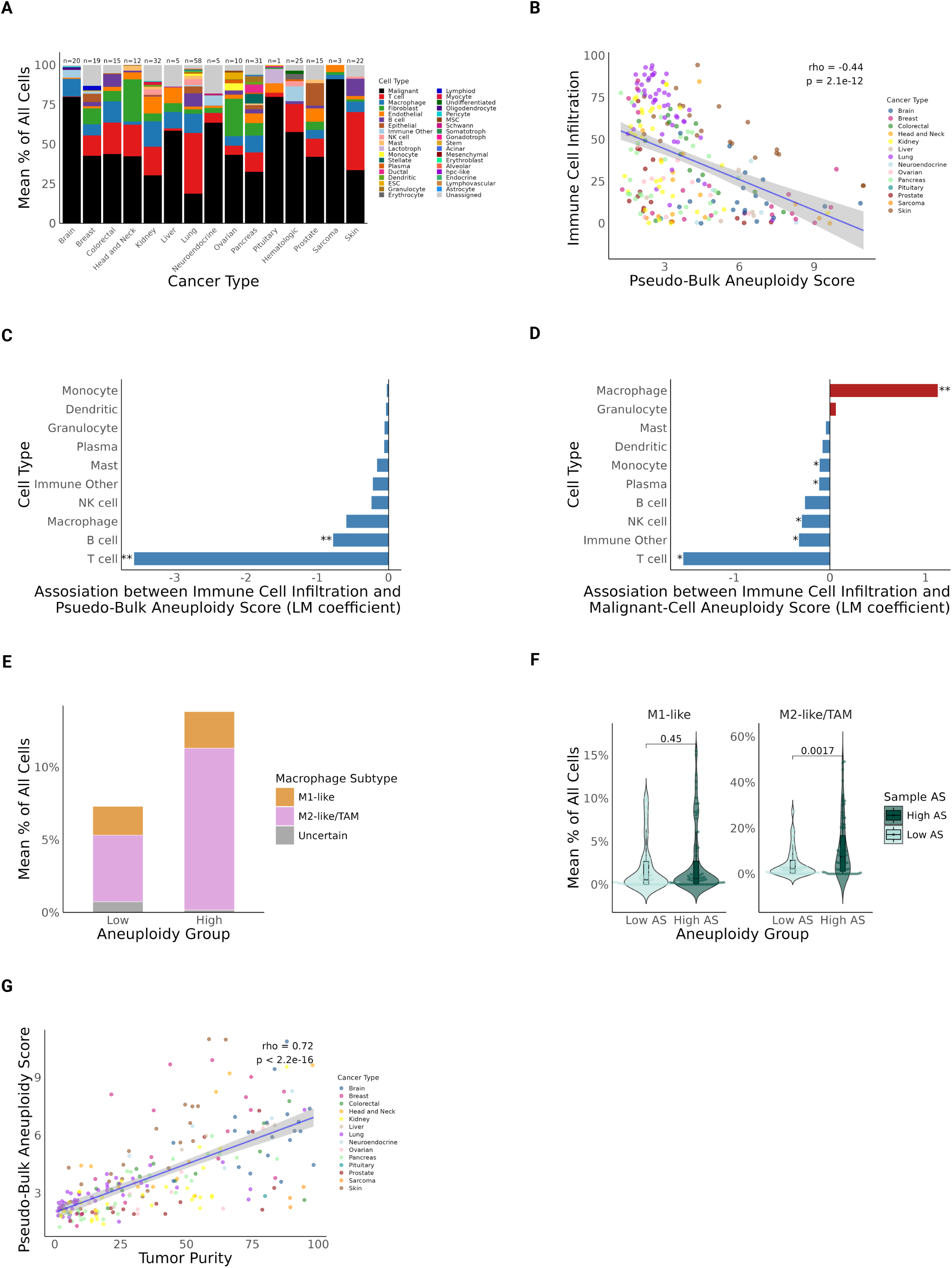
Tumor aneuploidy is associated with T-cell exclusion and infiltration of tumor-associated macrophages. **A)** Cell type abundance across cancer types: Stacked bar plot illustrating the mean relative abundances of malignant, immune, and stromal populations across 15 distinct cancer types (n = 273 tumor samples). Proportions reflect the percentage of all cells per sample, averaged within each cancer type. Sample size is annotated above each bar. Colors represent different cell types. **B)** Correlation between pseudo-bulk aneuploidy scores and immune infiltration: Scatter plot representing the relationship between the pseudo-bulk Aneuploidy Score (AS, x-axis) – calculated by averaging the AS across all cells in a sample – and immune cell abundance (y-axis). Each point represents a tumor (n = 248), color-coded by cancer type. A strong negative correlation is observed (Spearman’s rho = – 0.44, p = 2.1e–12), consistent with previous bulk-level observations. **C)** Associations of Immune cell types with pseudo-bulk AS: Bar plot displaying effect estimates (regression coefficients, x-axis) for specific immune cell types (y-axis) as a function of the pseudo-bulk AS. Coefficients were derived from linear regression models controlling for cancer-type-specific confounding effects. Blue and red bars indicate negative (depletion) and positive (enrichment) associations, respectively, with asterisks (*) denoting statistical significance (* q < 0.05, ** q < 0.01, *** q < 0.001). **D)** Association of immune cell types with malignant-cell-specific AS. Bars and asterisks as in (C). **E)** Mean relative abundance of macrophage phenotypes: Stacked bar plot illustrating the mean relative abundance of macrophage phenotypes (M1-like, M2-like/TAM, and Uncertain) calculated as a percentage of all cells within the sample. Tumors are stratified into discrete “Low” (bottom 30%) and “High” (top 30%) aneuploidy groups based on their malignant-cell AS. **F)** Distribution of macrophage subtypes by aneuploidy group: Violin plots detailing the distribution of M1-like and M2-like/TAM macrophage proportions across the High (dark green) and Low (light green) aneuploidy groups. Individual points represent tumor samples (two-sided Wilcoxon rank sum test, p = 0.45 for M1-like, 0.0017 for M2-like\TAM). **G)** Association between pseudo-bulk AS and tumor purity: Scatter plot comparing tumor purity (% malignant cells, x-axis) against the pseudo-bulk AS (y-axis). Each point represents a tumor (n = 248), color-coded by cancer type. A robust positive correlation is observed (Spearman’s rho = 0.72, p < 2.2e–16).

Previous tumor-bulk studies (3,4) demonstrated a downregulation of overall immune activity in highly aneuploid tumors. Consistent with these findings, evaluating the pseudo-bulk AS – calculated by averaging the AS across all cells in a sample – revealed a highly significant negative correlation with total immune infiltration (Fig. 5B). We used linear regression modeling to examine the association between aneuploidy and the abundance of various immune cell types, controlling for cancer type (Methods). All immune cell types were negatively associated with AS, with T and B cells being significantly depleted in aneuploid tumors (Fig. 5C and Supplementary Table S8). As traditional tumor-bulk analyses rely on computational inference, our approach provides a first direct single-cell-level confirmation of these previously reported associations.

Interestingly, however, when we substituted the pseudo-bulk metric with our malignant-cell-specific AS (Fig. 5D and Supplementary Table S8), we observed an interesting outlier. While T cells and nearly all other immune cell types remained depleted, macrophages (and, to a much lesser degree, granulocytes) became strongly and significantly enriched in the aneuploid tumors. These observations are concordant with a previous report in lung cancer, where induced aneuploidy was shown to drive macrophage and granulocyte accumulation in adjacent tissues while simultaneously suppressing T-cell infiltration (44). It is also consistent with the established link between aneuploidy, tumor aggressiveness, and disease progression (4,45): as tumors become more malignant, macrophages – and in particular Tumor-Associated Macrophages (TAMs) are continuously recruited to promote angiogenesis and invasion (46). Indeed, classification of the macrophage population into M1-like and M2-like/TAM subtypes (Supplementary Table S9-10) revealed that M2-like/TAMs were the most abundant subtype in highly aneuploid tumors (Fig. 5E). M2-like/TAMs – but not M1-like macrophages – were significantly more abundant in the High-AS vs (Fig. 5F). Furthermore, malignant-cell aneuploidy scores showed a significant positive association exclusively with the abundance of M2-like/TAM subtype (Supplementary Fig. S15A,B).

## Discussion

Our study leveraged a large pan-cancer scRNA-seq dataset (14) to explore the transcriptional consequences of aneuploidy at both cellular and tumor scales (Fig. 1). By jointly analyzing gene expression and inferred aneuploidy at the single-cell level, we observed notable differences in the transcriptional programs associated with high aneuploidy in single cells in comparison to bulk tumors. While tumor-level analyses have associated high aneuploidy with increased proliferation and metabolic activity (3,4), our single-cell comparisons revealed that highly aneuploid cells are often less proliferative, show reduced metabolic activity, and express more inflammatory-related signatures (Fig. 2). We show that these contrasting trends likely reflect differences in the timing and selection of karyotypic changes – whereas tumor-bulk data predominantly capture clonally selected, adapted aneuploidies, single-cell data detect emerging, potentially deleterious karyotypes subject to negative selection.

Our findings suggest that karyotypic heterogeneity plays an important role in shaping the cellular response to aneuploidy. Tumors with more heterogeneous aneuploidy profiles, containing a mixture of low-AS and high-AS cells, tend to exhibit reduced proliferation and metabolism in their most aneuploid subpopulations, alongside elevated p53 signaling and inflammatory pathways (Fig. 3). However, this is not the case in tumors with more clonally aneuploid profiles, likely reflecting adaptation and/or selection of more stable karyotypes. These patterns may help reconcile the so-called ‘aneuploidy paradox’ (2), by differentiating between the acute stress associated with newly acquired aneuploidy and the fitness-enhancing consequences of selected aneuploid states (47). A recent comprehensive mapping of gene expression changes following aneuploidy engineering *in vitro* found that emerging aneuploidy events drive p53-mediated stress and inflammatory signaling alongside a downregulation of cell cycle and metabolic programs (48). These cellular-level signatures appear to be specifically attenuated or lost in adapted, clonally stable cell populations (48). These experimental findings are highly consistent with our analysis of clinical tumor data.

We also investigated the transcriptional effects of recurrent chromosome-arm alterations across cancer types. While previous studies have suggested that copy number-defined genetic subclones often do not explain the majority of transcriptional heterogeneity (14), our results indicate that some recurrent aneuploidies are associated with consistent activation/repression of specific signaling pathways (Fig. 4). In some cases, these signatures could be associated with aneuploidy-induced changes in the expression of specific genes that reside on the gained/lost chromosome-arm. Notable examples include EMT upregulation in Chr9p-loss tumors, TGFβ signaling repression in Chr18q-loss tumors, and MYC pathway activation in Chr8q-gain tumors (32,37–40). These observations are consistent with the idea that some aneuploidies undergo positive selection due to their phenotypic impact on cellular programs relevant to tumor progression and adaptation.

Lastly, we explored at single-cell resolution the interactions between aneuploidy and the cell type composition of tumors. While our analysis recapitulated the well-established strong association between high degree of aneuploidy and reduced immune infiltration, we also found a strong and significant infiltration of macrophages into aneuploid tumors. We suspect that this association has been missed in previous tumor-bulk analyses due to the intrinsic association between tumor-bulk aneuploidy scores and tumor purity. Because tumor-bulk aneuploidy profiling aggregates signals from both malignant and non-malignant diploid cells, the resulting aneuploidy level is intrinsically sensitive to the proportion of malignant cells in the sample. Indeed, we observed a robust positive correlation between pseudo-bulk AS and tumor purity (defined as the fraction of malignant cells in the tumor; Fig. 5G), confirming that the pseudo-bulk metric is strongly affected by malignant cell content. Although prior studies employed computational methods to adjust for tumor purity (3,4), our results indicate that such adjustments may not fully decouple the aneuploidy signal from the underlying cell-type composition, emphasizing the importance of single-cell-level analyses.

Despite the resolution of our approach, we note the inherent limitations associated with inferring aneuploidy from transcriptomic data. A central challenge lies in balancing sensitivity and specificity when identifying copy number alterations. While prioritizing the reduction of false negatives can lead to the detection of false events in non-malignant control cells, applying stringent thresholds to minimize false positives inevitably results in the exclusion of true events in some malignant cells. Crucially, our findings remained robust across varying levels of filtering stringency. Nevertheless, future integration of single-cell DNA sequencing will be important. Furthermore, we observed associations between aneuploidy levels and immune cell infiltration at the tumor-level, but could not directly study the association between emerging aneuploid cells and their surrounding immune cells due to the difficulty to infer intercellular communication from dissociated single cells. Given the potential importance of such interactions for tumor evolution (4,49), it will be important to incorporate spatial transcriptomics and other methods to capture cell-cell interactions (50) to fully elucidate the role of aneuploidy in malignant-immune cell interactions.

In summary, our study offers a framework for investigating the functional consequences of aneuploidy in cancer by integrating high-resolution, single-cell transcriptomic and karyotypic data. By distinguishing between transient and selected aneuploid states, we reveal contrasting transcriptional profiles shaped by karyotypic heterogeneity and clonal selection. Additionally, we show that certain recurrent aneuploidies are associated with conserved gene expression changes, supporting their potential role in tumor adaptation. These findings contribute to a deeper understanding of how genome instability drives phenotypic diversity and may inform future strategies to exploit karyotypic features as biomarkers or therapeutic targets in cancer therapy.

## Methods

### Data Collection and Processing

Single-cell RNA sequencing (scRNA-seq) data were compiled from 304 tumor samples across 36 studies, spanning 15 distinct cancer types. This dataset was sourced from the Curated Cancer Cell Atlas (3CA, (14,18)) and included both gene expression profiles and related cell-level metadata. Gene-level copy number (CN) estimates for each cell were inferred using the InferCNA algorithm ((12), available at: [https://github.com/jlaffy/infercna]).

Data preprocessing followed the pipeline described by Gavish et al. (14). Briefly, cells with a low total UMI or TPM count were excluded from the analysis (for 10x data, the cutoff was >1,000 genes, and for SmartSeq2 data, the cutoff was >2,000 genes). Malignant cells were identified in each sample using the cell annotations provided in the original dataset. Samples with fewer than 10 malignant cells were excluded. Genes with low average expression across cells (log-transformed average TPM <4) were filtered out. Expression data were normalized using the transformation: log□(CPM / 10 + 1), followed by centering across genes. To ensure robust sample representation, only samples with >2,000 normalized expressed genes and >100 single cells with copy number data were retained. This pipeline resulted in matched GE and CN data for 234,744 malignant cells across 279 tumor samples.

### Chromosome-Arm Level Copy Number Calling

To obtain chromosome arm-level CN calls, gene-level CN matrices were processed using the ASCETS algorithm (51). Genomic arm coordinates were aligned to the hg38 reference, with a minimum arm breadth threshold set to 0.0. Chromosome-arm gains and losses were defined by a log□ ratio threshold of ±0.1, while all other parameters were kept at their default values.

### Aneuploidy Score (AS) Calculation

The Aneuploidy Score (AS) for each cell was calculated as the total number of chromosome-arms gained or lost, following the approach described by Cohen-Sharir et al. (19). The sample-level AS was defined as the mean AS across all malignant cells within each sample.

Cells were stratified into three aneuploidy categories: Low (AS ≤ 3), Mid (3 < AS < 7), and High (AS ≥ 7). Of the 279 samples, 114 samples included at least 30 malignant cells in both the Low and High AS groups and were therefore included in comparative downstream analyses.

### Differential Gene Expression and Pathway Enrichment Analyses

To evaluate the transcriptional consequences of aneuploidy at the single-sample level, we employed two complementary approaches for differential gene expression (DGE) analysis:

1. Group-based comparison: The *FindMarkers()* function from the Seurat R package was used to identify genes differentially expressed between High-AS and Low-AS malignant cell groups. A minimum detection threshold of 25% (min.pct = 0.25) was applied to include only genes expressed in at least 25% of cells in either group.
2. Continuous regression-based analysis: Gene-wise linear regression models were fitted within each sample, modeling gene expression as a function of AS (treated as a continuous variable across all malignant cells).

Following both analyses, pathway enrichment analysis was performed using the *enricher()* function from the clusterProfiler R package. Genes were selected for enrichment testing if they satisfied the following criteria: 1) log₂ fold change or regression coefficient ≠ 0, and 2) q < 0.25.

Significant genes were divided into upregulated and downregulated sets based on the direction of association, and sets with at least 10 significantly differentially expressed genes (q < 0.25) were used, enabling the identification of biological processes enriched in High-AS or Low-AS cells.

### Single-Sample GSEA and Meta-Analysis

To quantify pathway activity at the single-cell level, we performed single-sample Gene Set Enrichment Analysis (ssGSEA) using the *gsva()* function from the GSVA package with the ’ssgsea’ method. Enrichment scores were calculated for each pathway in each individual cell.

For each sample, t-tests were used to compare pathway enrichment between High-AS and Low-AS groups and Cohen’s d effect sizes were computed to quantify the magnitude of enrichment differences.

Subsequently, for each pathway, effect sizes and standard errors across samples were integrated using a random-effects meta-analysis, implemented with the *rma()* function in the metafor R package. This analysis generated a Pooled Effect Size reflecting the overall direction and magnitude of pathway enrichment differences across samples, and an FDR-adjusted q-value to assess statistical significance. This approach enabled robust inference of pathway-level trends associated with aneuploidy across multiple samples.

### Pseudo-Bulk Level Differential Gene Expression and Pathway Enrichment Analyses

For each of the 279 tumor samples, pseudo-bulk gene expression profiles were constructed by calculating the mean expression level of each gene across all tumor cells, using the log-transformed expression data prior to gene centering. Sample-level Aneuploidy Scores (AS) were computed as described above, based on arm-level copy number alterations.

Differential gene expression (DGE) analysis was conducted by fitting gene-wise linear regression models, where gene expression was modeled as a function of sample AS. To account for cancer-type-specific effects, tumor lineage was included as a covariate in each model.

Subsequent pathway enrichment analysis was performed using the same approach as described in the cell-level analysis above. Genes were retained for enrichment testing if they met both of the following criteria: 1) Regression coefficient ≠ 0, and 2) q < 0.25.

Significant genes were then divided into upregulated and downregulated sets based on the sign of the regression coefficient. Enrichment analysis was performed using the *enricher()* function from the clusterProfiler R package to identify biological pathways associated with higher or lower AS.

The same analytical pipeline was applied to two additional pseudo-bulk settings:

1. gene expression aggregated only across malignant cells, and
2. a subset of samples that contained at least 30 cells in both the Low and High AS groups.

Additionally, an alternative DGE analysis was performed using all 279 tumor samples, with both tumor lineage and tumor purity included as covariates in each gene-wise linear regression model. Tumor purity was defined as the percentage of malignant cells in the tumor.

### TCGA Tumo-Bulk Differential Gene Expression and Pathway Enrichment Analyses

For the TCGA dataset, we utilized the ranked list of genes obtained from a linear regression model performed by Taylor et al. (3), in which gene expression was regressed on aneuploidy score (AS) while controlling for tumor type as a covariate to adjust for inter-cancer variability. This model generated gene-specific regression coefficients, representing the strength and direction of association between AS and gene expression across diverse tumor types.

Following this, pathway enrichment analysis was conducted following the same methodology as in the cell-level and pseudo-bulk analyses described above. Genes were included if they met the following criteria: 1) Regression coefficient ≠ 0, and 2) Bonferroni-adjusted p-value < 0.25.

Genes were stratified into upregulated and downregulated categories based on the direction of the coefficient, and functional enrichment was assessed using the *enricher()* function from the clusterProfiler package to identify pathways linked to increased or decreased aneuploidy.

This analysis was also performed using a ranked list of genes obtained from Taylor et al., in which gene expression was regressed against AS while controlling for both tumor type and tumor purity as covariates.

Additionally, gene set enrichment analysis (GSEA) was performed using the ranked list of genes obtained from Taylor et al., with the *GSEA()* function from the clusterProfiler R package set to default parameters.

### Quality Control and Robustness Analyses

Cellular fitness and sequencing quality were assessed using the number of expressed genes and the mitochondrial transcript fraction per cell. Based on initial distributions, cells with <1,500 expressed genes or a mitochondrial fraction >10% were excluded. Single-cell DGE and pathway enrichment analyses were re-conducted on this subset of cells (n = 113,142 malignant cells across 81 tumor samples).

To ensure findings were independent of aneuploidy inference parameters, a sensitivity analysis was performed using more stringent scoring criteria for the ASCETS algorithm. (51). For this analysis, the minimum chromosome-arm breadth threshold was set to 0.1, and chromosome-arm gains and losses were defined by a log₂ ratio threshold of ±0.2, with all other parameters kept at default values. These conservative parameters resulted in a downward shift of the AS distribution, and group thresholds were consequently adjusted to: Low (AS ≤ 1), Mid (1 < AS < 4), and High (AS ≥ 4). Downstream comparative analyses, including single-cell ssGSEA meta-analysis and pseudo-bulk DGE and pathway enrichment analyses, were restricted to 104 tumors retaining ≥30 malignant cells in both the Low-AS and High-AS groups.

Finally, to address potential confounding effects of cell cycle distribution on copy number inference and gene expression, cells were stratified by cell cycle phase (G1, S, and G2/M) using the *CellCycleScoring()* function in the Seurat R package. The gene sets used for scoring were the canonical S and G2/M marker lists defined by Gavish et al. (14). Single-cell ssGSEA meta-analysis and pseudo-bulk DGE and enrichment analyses were performed separately for non-cycling (G1) and cycling (S/G2/M) populations. To ensure robust statistical power and stable mean estimates, these analyses were restricted to tumor samples retaining a minimum of 100 malignant cells within the relevant cell cycle phases. Specifically, the non-cycling analysis included 173 tumor samples (70 of which met the 30-cell threshold per AS group), while the cycling analysis included 181 tumor samples (73 of which met the 30-cell threshold per AS group).

### Intratumoral Aneuploidy Heterogeneity Calculation

To quantify intra-tumor aneuploidy heterogeneity scores, pairwise cell-to-cell similarity scores were computed within each sample based on gene-level copy number profiles of malignant cells. Specifically, Pearson’s correlation coefficients were calculated between all pairs of malignant cells in a sample, using their inferred gene-level copy number vectors.

For each sample, the mean of all pairwise correlations was computed to define a Karyotypic Heterogeneity Score. This metric reflects the overall similarity in copy number profiles among malignant cells.

To investigate the influence of intra-tumor karyotypic heterogeneity on pathway-level alterations, we repeated the random-effects meta-analysis described previously, this time stratifying samples based on their karyotypic heterogeneity scores. Tumors were divided into tertiles, and separate meta-analyses were conducted for the top one-third (most heterogeneous tumors, n=43) and bottom one-third (most homogeneous tumors, n=24).

### Copy Number Subclone Identification and Merging

To identify copy number-defined subclones and quantify subclonal heterogeneity within individual tumors, we applied a Louvain clustering approach using the Seurat framework on filtered copy number alteration (CNA) matrices. Each CNA matrix was pre-processed by retaining the top 67% of genes with the highest absolute CNA values.

Subclones were inferred using Seurat’s clustering pipeline. A shared nearest neighbor (SNN) graph was constructed using the *FindNeighbors()* function with k.param = 15 and dims = 1:15. Louvain clustering was then performed using the *FindClusters()* function with a resolution parameter of 0.5.

To refine and consolidate subclonal boundaries, a post-clustering merging step was implemented following criteria similar to those described by Gavish et al. (14). Specifically, for each chromosomal arm, the average CNA (Copy Number Alteration) value across genes on that arm was computed for each cluster. A chromosomal arm was considered amplified within a cluster if the average CNA exceeded 0.15 or deleted if it fell below −0.15. Chromosomal arms with average CNA values between −0.15 and 0.15 were classified as neutral. Clusters that exhibited identical chromosomal arm-level CNA profiles (i.e., all chromosome arms had the same status) or where the maximum difference in average CNA values between arms within a cluster was less than 0.15 were iteratively merged. This process resulted in a final set of genetically distinguishable subclones for each tumor.

Next, DGE and pathway enrichment analyses were performed to assess transcriptional differences between subclones. For each tumor, individual subclones were compared against all other subclones from the same tumor using the *FindMarkers()* function in Seurat, employing the Wilcoxon rank-sum test and a detection threshold of 25% (min.pct = 0.25). Genes with a log₂ fold change ≠ 0 and q < 0.25 were retained for enrichment analysis. Genes were stratified into upregulated and downregulated sets based on the direction of the fold change, and only sets with at least 10 significant genes were considered. Pathway enrichment analysis was conducted using the *enricher()* function from the clusterProfiler R package to identify biological processes significantly enriched or depleted in each subclone.

### Recurrent Chromosome-Arm-Level Alterations and Transcriptomic Analysis of TCGA data

To identify recurrent chromosome-arm-level alterations across human cancers, we utilized the Cancer Genome Atlas (TCGA) cohort, following the methodology described in Saad et al. (17). Gene expression data (FPKM) were retrieved using the TCGAbiolinks R package. Arm-level copy number status and aneuploidy scores for each sample were obtained from Supplementary Table S2 of Taylor et al. (3). The dataset included approximately 7,500 primary tumor samples spanning more than 20 cancer types. An arm-level gain or loss was defined as common in a given cancer type if it occurred in more than 20% of tumor samples for that type.

This analysis identified a total of 230 common arm losses and 196 common arm gains across the TCGA cohort. For recurrent arm losses, previously published GSEA results from Saad et al. (17) were used. For recurrent arm gains, we conducted a new differential gene expression (DGE) and enrichment analysis using the exact same methodology. To account for the overall aneuploidy levels as a potential confounder, DGE was performed using inverse probability of treatment weighting (IPTW). Weights were derived from each sample’s aneuploidy score. For each gene, a weighted version of Cohen’s *d* statistic was computed to estimate log fold-changes between arm-gained and non-arm-gained tumors, applied to log□□-transformed FPKM expression values, and incorporating the sample-specific weights. Following DGE, gene set enrichment analysis (GSEA) was performed using the *GSEA()* function from the clusterProfiler R package with default parameters. Enrichment testing was carried out using the ranked list of differentially expressed genes for each recurrent chromosome-arm gain, evaluating both ‘Hallmark’ and ‘Reactome’ gene sets to identify biological processes associated with recurrent copy number gains across cancer types.

### Recurrent Chromosome-Arm-Level Alterations and Transcriptomic Analysis of Single-Cell Data

To investigate recurrent chromosome-arm-level alterations and their transcriptional consequences at single-cell resolution in our cohort of 279 tumor samples, recurrent chromosome arm–cancer type (CA–CT) pairs were defined, as described above. For each CA–CT pair, cells from each tumor sample were divided into two groups: those with and those without the specific recurrent aneuploidy. CA–CT pairs were included in the analysis only if they had at least 5 samples, and for each sample, both cell groups comprised >30 cells. This filtering resulted in 57 CA–CT pairs for recurrent arm losses (across 7 cancer types) and 25 CA–CT pairs for recurrent arm gains (across 6 cancer types).

To assess pathway activity at the single-cell level, we performed ssGSEA using the *gsva()* function from the GSVA R package with the ’ssgsea’ method. Enrichment scores were computed for each cell across both Hallmark and Reactome gene sets. Within each sample, a t-test was used to compare ssGSEA enrichment scores between cells with and without the recurrent chromosome-arm alteration, and the effect size was calculated to quantify the enrichment differences between the two cell groups.

To integrate findings across samples, we performed a random-effects meta-analysis using the *rma()* function from the metafor R package. This approach generated, for each pathway, a pooled effect size representing the average magnitude and direction of enrichment difference across samples, along with a corresponding false discovery rate (FDR)-adjusted q-value to assess statistical significance. In parallel with the ssGSEA-based meta-analysis, DGE and GSEA analyses were also performed at the single-cell level using the same IPTW-based framework applied to the TCGA cohort above. Cell-specific weights, derived from each cell’s aneuploidy score, were used to compute a weighted version of Cohen’s d statistic, comparing normalized expression values between cells with and without the arm alteration. Genes were then ranked and tested for pathway enrichment using the *GSEA()* function from the clusterProfiler R package (default settings), evaluating both Hallmark and Reactome gene sets.

### Correlation Between TCGA and Single-Cell Pathway Enrichment Results

To assess the agreement between bulk and single-cell analyses, the correlations between pathway enrichment results were calculated across matched chromosome-arm–cancer type (CA–CT) pairs for which both TCGA and single-cell datasets were available.

For each such CA–CT pair, normalized enrichment scores (NES) were extracted from the TCGA bulk analysis, and pooled effect sizes were extracted from the single-cell meta-analysis, for both ‘Hallmark’ and ‘Reactome’ gene sets. The analysis was restricted to pathways that were significantly enriched in both datasets (q < 0.25). Spearman’s correlation coefficients were then computed between TCGA NES values and single-cell pooled effect sizes to quantify the consistency of pathway-level responses to recurrent chromosome-arm alterations.

### Analysis of Tumor Microenvironment Composition and Immune-Aneuploidy Interactions

Cell-type abundances were obtained from the 3CA (14,18), quantified for each sample, and expressed as a percentage of total cells. Tumor purity was defined as the malignant cell fraction within each sample. The total immune cell fraction was calculated by aggregating the abundances of T cells, B cells, macrophages, natural killer (NK) cells, mast cells, monocytes, dendritic cells (including pDCs), plasma cells, and granulocytes. Cells with lower-resolution annotations – specifically broad categories such as myeloid, lymphoid, and tumor-infiltrating lymphocytes (TILs) – were grouped as “Other immune” and included in the total immune calculation. Any cells lacking clear annotations (e.g., “Other”, “Unknown”, or “non-malignant”) were categorized as “Unassigned”. Samples were excluded from downstream analysis if the “Unassigned” category exceeded 40% of their total cell count (n=6) or if they were derived from hematological malignancies (n=25).

Pan-cancer associations between AS and immune infiltration, or between tumor purity and immune infiltration, were assessed using two-sided Spearman’s correlations. Pseudo-bulk AS was computed by averaging the AS across all cells in a sample. To account for cancer-type-specific confounding variables across the cohort, we utilized linear regression models for each individual immune population. These models evaluated immune cell fractions as a function of aneuploidy score while controlling for tumor lineage as a covariate.

Regression coefficients and their associated p-values were used to determine the independent effect of aneuploidy on specific immune subsets. This analysis was performed using two distinct metrics: 1) the pseudo-bulk AS and 2) the malignant-cell-specific AS.

For high-resolution characterization of the myeloid compartment, macrophages were isolated based on global annotations and integrated using the Harmony algorithm across the first 30 principal components to mitigate tissue-of-origin batch effects. Unsupervised clustering was performed using the Louvain algorithm (resolution 0.3) on a Shared Nearest Neighbor (SNN) graph derived from the first 20 Harmony-corrected dimensions. Clusters were assigned biological identities independently of the marker genes used for subsequent testing using the *AddModuleScore()* function from the Seurat R package (Supplementary Table S9). This function evaluated the mean expression of curated canonical gene sets representing M1-like, M2-like, and specific TAM subpopulations (C1Q, SPP1, ISG15, FN1), adapted from Azizi et al. (52) and Cheng et al. (53). Clusters were designated as “Uncertain” if no gene set achieved a mean score greater than zero; for broader analyses, TAM subpopulations and M2-like clusters were unified into a single “TAM/M2-like” category.

To evaluate subtype-specific enrichment, samples were stratified into High (top 30%) and Low (bottom 30%) aneuploidy groups based on their malignant-cell AS. Proportional differences between these groups were evaluated using Wilcoxon rank-sum tests. To model the independent effect of continuous aneuploidy while controlling for cancer lineage, linear regression models were applied for each subtype.

## Statistical Analyses

All computational and statistical analyses were conducted in R (v4.3.0 and v4.3.1). Differential gene expression (DGE) was evaluated using two-sided Wilcoxon rank-sum tests for group-based comparisons or gene-wise linear regression for continuous analysis, incorporating tumor lineage and purity as covariates where indicated. Functional enrichment was assessed using two complementary frameworks: (1) over-representation analysis of significantly differentially expressed genes via the *enricher()* function, and (2) single-sample Gene Set Enrichment Analysis (ssGSEA). For the latter, pathway-level differences between groups were first evaluated within individual samples using two-sided Welch’s t-tests and Cohen’s d effect sizes, which were subsequently integrated across the cohort via a random-effects meta-analysis model. All p-values were adjusted for multiple hypothesis testing using the Benjamini–Hochberg False Discovery Rate (FDR) procedure, except for TCGA analyses, which utilized Bonferroni adjustment. Statistical significance was defined at a discovery threshold of q < 0.25 for exploratory pathway enrichment and a stringent q < 0.05 for immune infiltration analysis. Relationships between continuous variables were quantified using Spearman’s correlation coefficients.

## Data Availability

Single-cell RNA-seq data were obtained from the 3CA (14,18). Publicly available TCGA data were accessed via the TCGAbiolinks R package. Recurrent chromosome-arm losses and corresponding pathway enrichment results were obtained from Saad et al. (17). Chromosome-arm copy number calls, recurrent chromosome-arm gains, and pathway enrichment results are included within the article as Supplementary Data.

## Authors’ Contributions

**G. Wolf-Dankovich** designed and performed the bioinformatic analyses, interpreted results, generated figures, and wrote the manuscript. **T. Mashiah** designed and executed the immune microenvironment and cell-type abundance analyses and edited the manuscript. **R. Saad** and **H. Khoury** contributed to code development and analysis support. **I. Tirosh** and **E. Somech** generated the 3CA dataset, assisted with data curation, generated the single-cell gene-level copy number data from the scRNAseq data, and edited the manuscript. **U. Ben-David** conceived, supervised, and directed the study, interpreted results, and wrote the manuscript.

## Acknowledgments

We thank the members of the Ben-David laboratory for helpful discussions. Work in the Ben-David laboratory is supported by the Israel Cancer Research Fund Project Award, the Israel Science Foundation project grant (1805/21), the U.S.-Israel Binational Science Foundation (BSF) project grant (2019228), the European Research Council Starting Grant (945674), and the Schmidt Science Polymaths Award. Artificial intelligence was used for language editing.

## Supplementary Tables

**Supplementary Table 1: Chromosome-Arm Level Copy Number Profiles for 496,467 single cells.**

Cell-specific chromosome-arm level copy number estimates for all samples, as determined by the ASCETS algorithm (n=496,467).

**Supplementary Table 2: Single-Sample GSEA and Meta-Analysis Results.** Pooled effect sizes and statistical significance for ‘Hallmark’ gene sets, derived from analyses of all tumors and after stratification based on karyotypic heterogeneity scores.

**Supplementary Table 3: Pseudo-Bulk Differential Gene Expression and Pathway Enrichment Analysis.**

Pathway enrichment results for ‘Hallmark’ gene sets, identifying pathways associated with higher or lower AS. The analysis was performed using gene-wise linear regression models, adjusting for tumor lineage and the combination of tumor lineage and tumor purity.

**Supplementary Table 4: Recurrent Chromosome-Arm Gains Across the TCGA Cohort.**

Table detailing recurrent chromosome arm gains observed in the TCGA cohort.

**Supplementary Table 5: Gene Set Enrichment Analysis (GSEA) Results for Recurrent Chromosome-Arm Alterations in Single-Cell Data.**

GSEA results for ‘Hallmark’ and ‘Reactome’ gene sets, analyzing recurrent chromosome-arm-level losses and gains in single-cell data.

**Supplementary Table 6: Gene Set Enrichment Analysis (GSEA) Results for Recurrent Chromosome Arm-Gains in the TCGA Cohort.**

GSEA results for ‘Hallmark’ and ‘Reactome’ gene sets, analyzing recurrent chromosome-arm-level gains in the TCGA cohort.

**Supplementary Table 7: Cell Type Abundance Across Cancer Types.**

Cell-type abundances across the cohort, quantified as the percentage of total cells for each sample and stratified by cancer type.

**Supplementary Table 8: Regression Analysis of Immune Infiltration and Aneuploidy.**

Linear regression results evaluating the association between immune cell fractions and both pseudo-bulk AS and malignant-cell AS, and between macrophage subtypes and malignant-cell AS, while controlling for tumor lineage as a covariate.

**Supplementary Table 9: Macrophage Sub-cluster Annotation and Module Scoring Statistics.**

Annotation of macrophage clusters based on canonical gene sets (M1-like, M2-like, and specific TAM subpopulations). Biological identities were assigned using the *AddModuleScore()* function from the Seurat R package to evaluate the mean expression of curated signatures.

**Supplementary Table 10: Sample-Level Macrophage Subtype Proportions and Aneuploidy Stratification.**

Table summarizing the cell counts and proportional abundances of macrophage subtypes within individual tumor samples. The table details sample-specific cell counts (macrophage subtypes and total cells) and their corresponding stratification into “High,” “Low,” or “Intermediate” aneuploidy groups based on cancer-type-specific quintile thresholds.

