## Supplementary Figures for "Pan-cancer analysis of single-cell RNA sequencing data from 304 human tumors sheds light on the ‘aneuploidy paradox’"

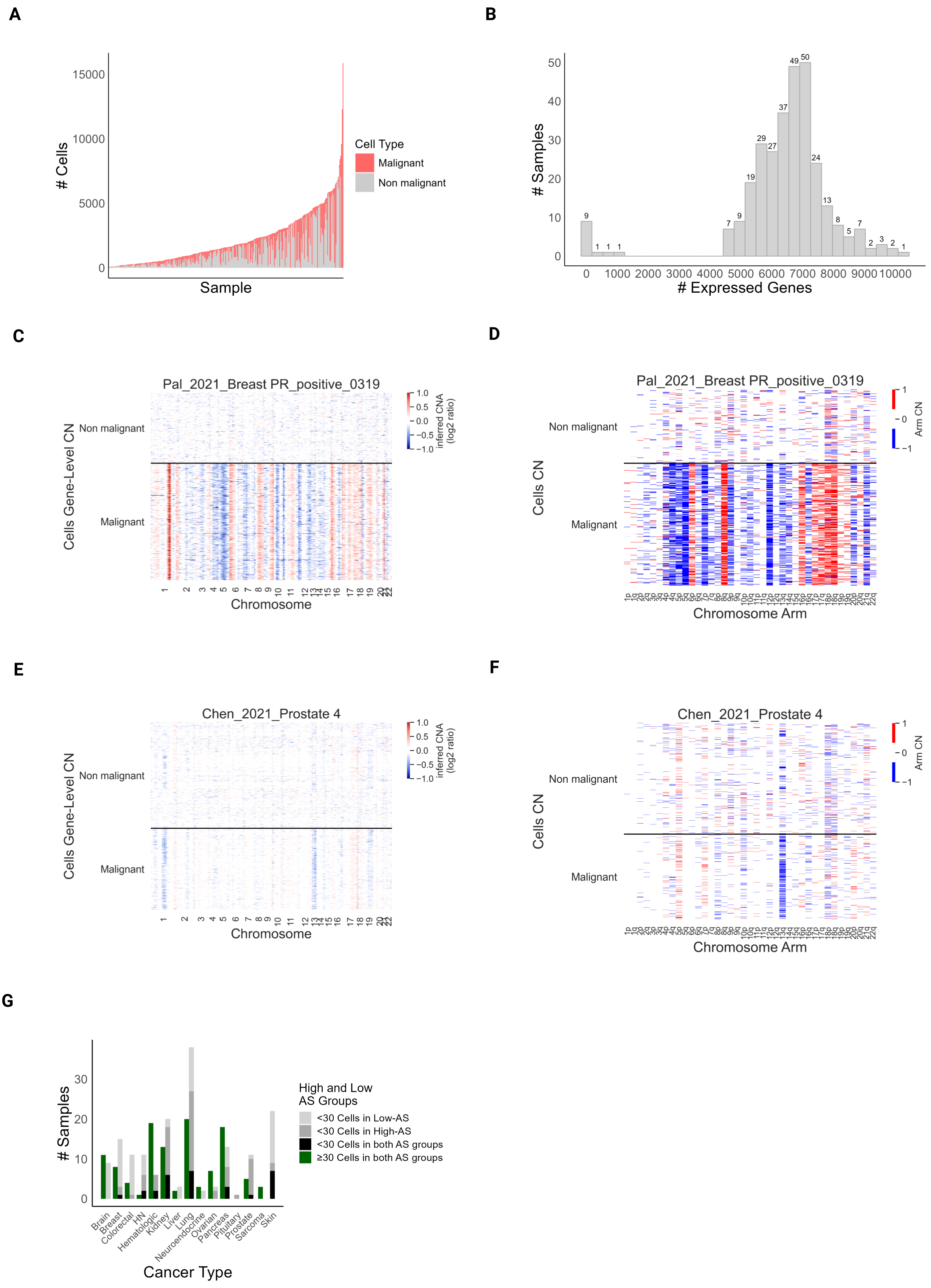
A
B
C
D
E
F
G
### Cells
Sample
Cell Type
Malignant
Non malignant
### Samples
### Expressed Genes
Pal_2021_Breast PR_positive_0319
Cells Gene-Level CN
Non malignant
Malignant
Chromosome
inferred CNA (log2 ratio)
Cells CN
Chromosome Arm
Arm CN
Chen_2021_Prostate 4
Cancer Type
High and Low AS Groups
<30 Cells in Low-AS
<30 Cells in High-AS
<30 Cells in both AS groups
≥30 Cells in both AS groups
Brain
Breast
Colorectal
HN
Hematologic
Kidney
Liver
Lung
Neuroendocrine
Ovarian
Pancreas
Pituitary
Prostate
Sarcoma
Skin

#### Supplementary Figure S1: Tumor composition and pipeline demonstration (related to Fig. 1)

**A)** Cell type distribution across tumor samples: Stacked bar plot showing the number of cells per tumor (total n = 665,842 cells across 304 tumor samples). The X-axis represents individual tumors, and the Y-axis indicates cell count. Bars are color-coded to distinguish malignant (red) from non-malignant (gray) cells. **B)** Distribution of the total number of genes expressed per tumor: A histogram showing the number of tumors based on gene expression. The X-axis shows the total number of genes expressed per tumor (a total of 1,954,122 genes across 304 tumors), while the Y-axis indicates the number of tumor samples that fall into each bin. Tumors with low gene counts (less than 2000) were excluded from downstream analysis. **C)** Copy number profiles of single cells from a representative highly aneuploid tumor: Heatmaps showing gene-level copy number (CN) alterations in a representative highly aneuploid tumor. Each column represents a gene; each row represents a single tumor cell. Color indicates inferred CNA (log2 ratio) values (blue = loss; red = gain). **D)** Arm-level CN profiles of the same representative highly aneuploid tumor: Heatmaps showing arm-level CN alterations. Each column represents a chromosome arm; each row represents a single tumor cell. Color indicates arm CN (blue = loss; red = gain). **E)** Copy number profiles of single cells from a representative near-diploid karyotype tumor: Heatmaps as in (C). **F)** Arm-level CN profiles of the same representative near-diploid karyotype tumor: Heatmaps as in (D). **G)** AS group-based tumor composition across cancer types: Bar plot showing the number of tumors per cancer type (n = 15), stratified by whether they contain both High-AS and Low-AS cell groups. Green bars indicate the number of tumors that include both high-AS and low-AS cells per cancer type (n=114); gray bars represent tumors that do not include ≥30 cells from both groups.

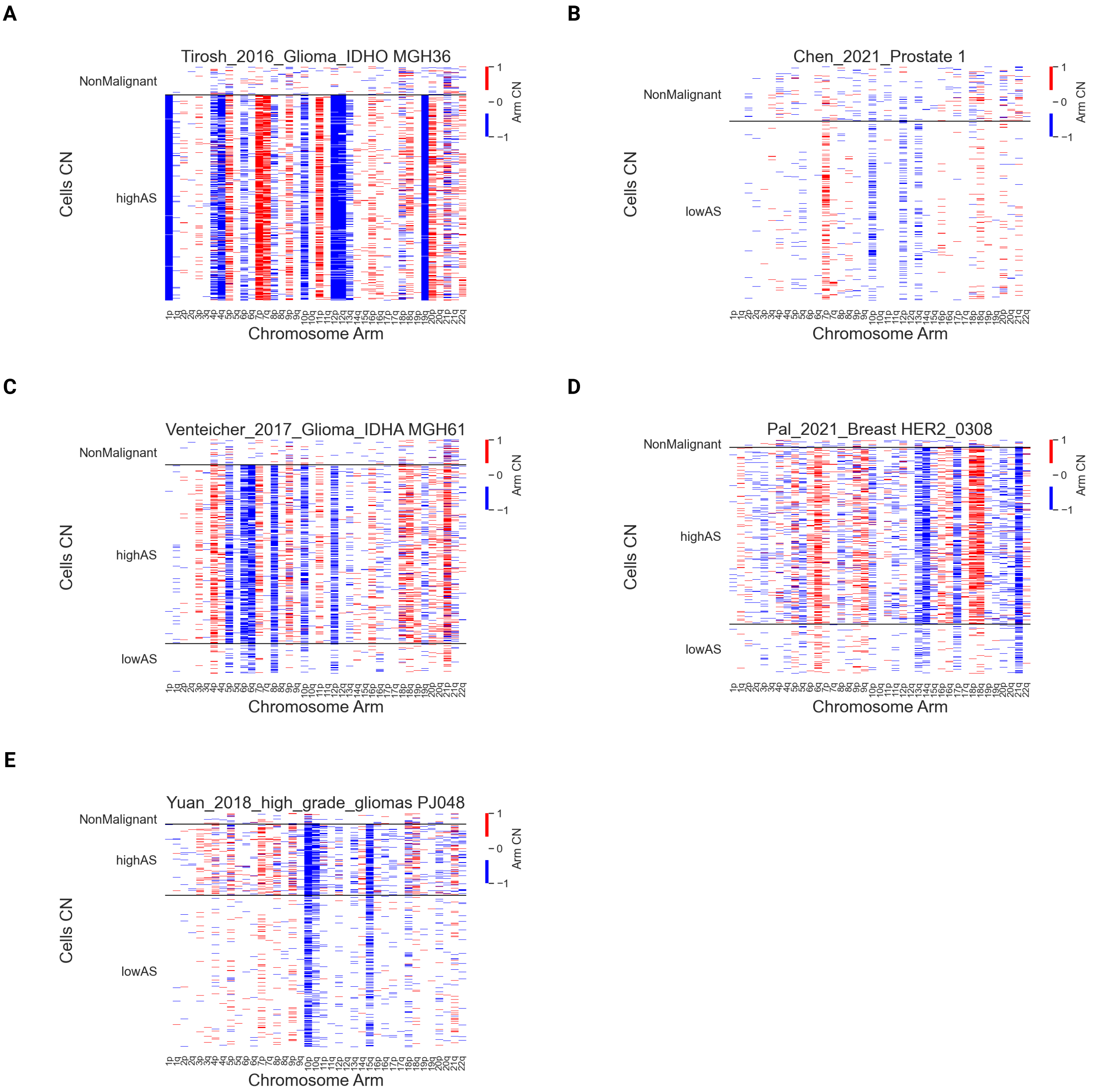
A
Tirosh_2016_Glioma_IDHO MGH36
NonMalignant
highAS
Cells CN
Chromosome Arm
Arm CN
1
0
−1
B
Chen_2021_Prostate 1
NonMalignant
lowAS
Cells CN
Chromosome Arm
Arm CN
C
Venteicher_2017_Glioma_IDHA MGH61
NonMalignant
highAS
lowAS
Cells CN
Chromosome Arm
Arm CN
D
Pal_2021_Breast HER2_0308
NonMalignant
highAS
lowAS
Cells CN
Chromosome Arm
Arm CN
E
Yuan_2018_high_grade_gliomas PJ048
NonMalignant
highAS
lowAS
Cells CN
Chromosome Arm
Arm CN

#### **Supplementary Figure S2: Inferred arm-level aneuploidy profiles (related to Fig. 1)**

**A-E)** Arm-level copy number profiles across diverse tumor contexts: Heatmaps showing chromosomal arm-level copy number (CN) alterations in representative tumors, including (A) highly aneuploid, (B) near-diploid, and (C-E) heterogeneous tumors containing a mixture of Low-AS and High-AS malignant cells. Each column represents a chromosomal arm; each row represents a single cell. Cells are grouped by their classification as non-malignant or malignant (Low-AS, Mid-AS, or High-AS). Color indicates inferred arm-level CN values (red = gain; blue = loss)

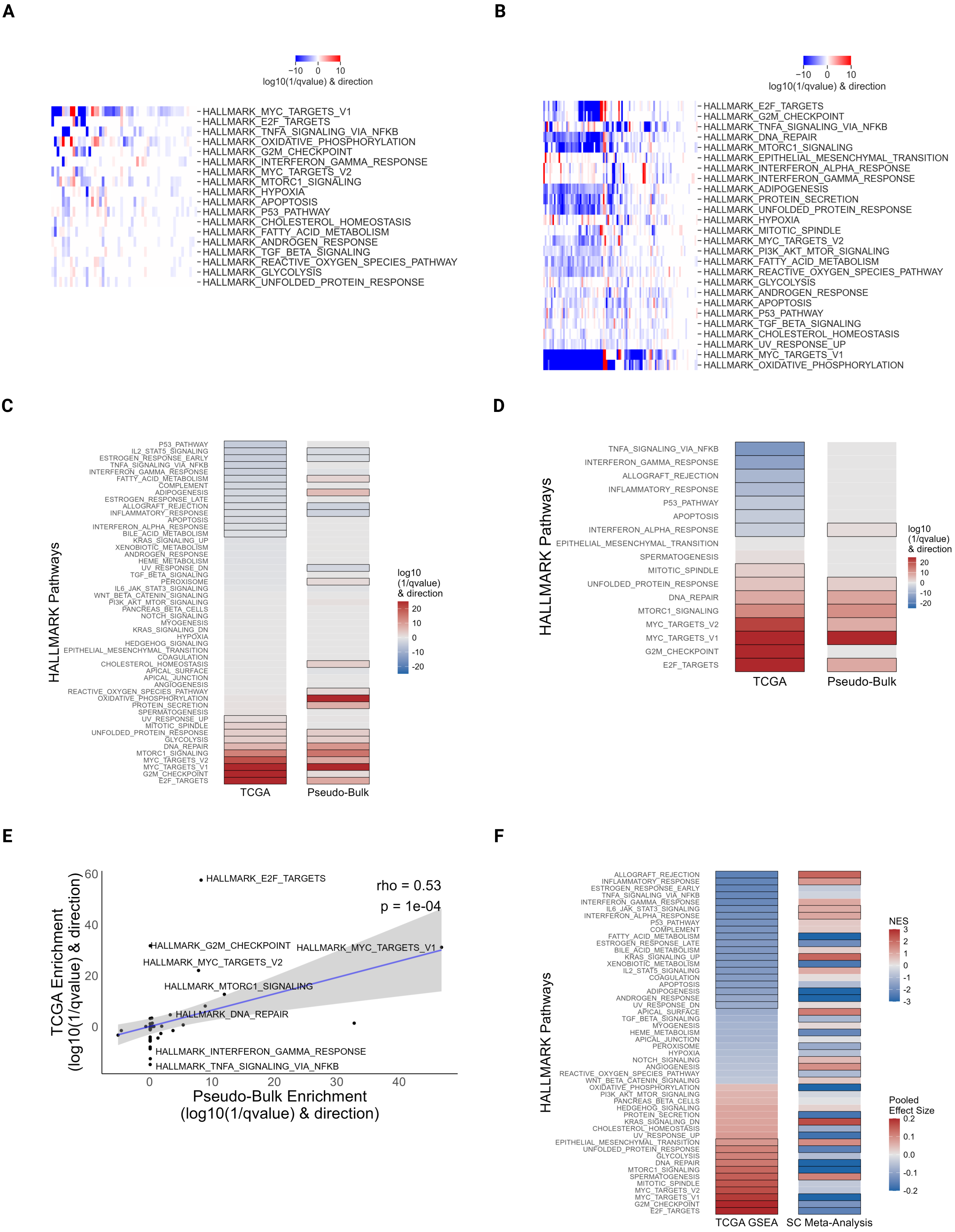
A
B
−10 0 10
log10(1/qvalue) & direction
HALLMARK_MYC_TARGETS_V1
HALLMARK_E2F_TARGETS
HALLMARK_TNFA_SIGNALING_VIA_NFKB
HALLMARK_OXIDATIVE_PHOSPHORYLATION
HALLMARK_G2M_CHECKPOINT
HALLMARK_INTERFERON_GAMMA_RESPONSE
HALLMARK_MYC_TARGETS_V2
HALLMARK_MTORC1_SIGNALING
HALLMARK_HYPOXIA
HALLMARK_APOPTOSIS
HALLMARK_P53_PATHWAY
HALLMARK_CHOLESTEROL_HOMEOSTASIS
HALLMARK_FATTY_ACID_METABOLISM
HALLMARK_ANDROGEN_RESPONSE
HALLMARK_TGF_BETA_SIGNALING
HALLMARK_REACTIVE_OXYGEN_SPECIES_PATHWAY
HALLMARK_GLYCOLYSIS
HALLMARK_UNFOLDED_PROTEIN_RESPONSE
HALLMARK_E2F_TARGETS
HALLMARK_G2M_CHECKPOINT
HALLMARK_TNFA_SIGNALING_VIA_NFKB
HALLMARK_DNA_REPAIR
HALLMARK_MTORC1_SIGNALING
HALLMARK_EPITHELIAL_MESENCHYMAL_TRANSITION
HALLMARK_INTERFERON_ALPHA_RESPONSE
HALLMARK_INTERFERON_GAMMA_RESPONSE
HALLMARK_ADIPOGENESIS
HALLMARK_PROTEIN_SECRETION
HALLMARK_UNFOLDED_PROTEIN_RESPONSE
HALLMARK_HYPOXIA
HALLMARK_MITOTIC_SPINDLE
HALLMARK_MYC_TARGETS_V2
HALLMARK_PI3K_AKT_MTOR_SIGNALING
HALLMARK_FATTY_ACID_METABOLISM
HALLMARK_REACTIVE_OXYGEN_SPECIES_PATHWAY
HALLMARK_GLYCOLYSIS
HALLMARK_ANDROGEN_RESPONSE
HALLMARK_APOPTOSIS
HALLMARK_P53_PATHWAY
HALLMARK_TGF_BETA_SIGNALING
HALLMARK_CHOLESTEROL_HOMEOSTASIS
HALLMARK_UV_RESPONSE_UP
HALLMARK_MYC_TARGETS_V1
HALLMARK_OXIDATIVE_PHOSPHORYLATION
C
D
HALLMARK Pathways
log10 (1/qvalue) & direction
20
10
0
-10
-20
TCGA
Pseudo-Bulk
TNFA_SIGNALING_VIA_NFKB
INTERFERON_GAMMA_RESPONSE
ALLOGRAFT_REJECTION
INFLAMMATORY_RESPONSE
P53_PATHWAY
APOPTOSIS
INTERFERON_ALPHA_RESPONSE
EPITHELIAL_MESENCHYMAL_TRANSITION
SPERMATOGENESIS
MITOTIC_SPINDLE
UNFOLDED_PROTEIN_RESPONSE
DNA_REPAIR
MTORC1_SIGNALING
MYC_TARGETS_V2
MYC_TARGETS_V1
G2M_CHECKPOINT
E2F_TARGETS
E
F
60
40
20
0
0 10 20 30 40
TCGA Enrichment (log10(1/qvalue) & direction)
Pseudo-Bulk Enrichment (log10(1/qvalue) & direction)
rho = 0.53
p = 1e-04
HALLMARK_E2F_TARGETS
HALLMARK_G2M_CHECKPOINT
HALLMARK_MYC_TARGETS_V1
HALLMARK_MYC_TARGETS_V2
HALLMARK_MTORC1_SIGNALING
HALLMARK_DNA_REPAIR
HALLMARK_INTERFERON_GAMMA_RESPONSE
HALLMARK_TNFA_SIGNALING_VIA_NFKB
NES
3
2
1
0
-1
-2
-3
Pooled Effect Size
0.2
0.1
0.0
-0.1
-0.2
TCGA GSEA
SC Meta-Analysis

**Supplementary Figure S3: Expanded gene set analyses highlight intra-tumor trends and confirm the inverse relationship between single-cell and bulk cell-population data (related to Fig. 2)**

**A)** Intra-tumor pathway enrichment in single cells: Heatmap showing differential enrichment of 'Hallmark' gene sets between High-AS and Low-AS cells within individual tumors. Each column represents a tumor (n=55). Colors indicate direction and strength of enrichment (red = upregulated; blue = downregulated in High-AS cells). **B)** Intra-tumor pathway enrichment using a linear model: Heatmap showing pathway enrichment based on a linear regression model assessing the association between AS and pathway scores within each tumor (n=101). Columns represent tumors; Colors indicate direction and strength of enrichment (red = upregulated; blue = downregulated in High-AS cells). **C)** Inter-tumor pathway enrichment in bulk tumors: Enrichment analysis comparing High-AS vs. Low-AS tumors in both pseudo-bulk (aggregated from single-cell data) and TCGA datasets across all 'Hallmark' gene sets. Heatmap colors represent enrichment direction and strength (red = upregulated; blue = downregulated in High-AS). Black boxes indicate significant pathways ($q < 0.05$). **D)** Bulk tumor enrichment analysis controlling for tumor purity: Enrichment analysis comparing High-AS vs. Low-AS tumors in both TCGA and pseudo-bulk datasets, with tumor purity included as a covariate. Shown are selected Hallmark pathways related to proliferation, metabolism, and immunity. Colors represent direction and strength of enrichment; black boxes highlight significant pathways ($q < 0.05$). **E)** Correlation of pathway changes across bulk datasets with purity control: Scatter plot showing the correlation between pseudo-bulk and TCGA enrichment scores when controlling for tumor purity. The x-axis shows pseudo-bulk enrichment values; the y-axis shows TCGA enrichment values. A significant positive correlation is observed (Spearman's rho = 0.53, p = 1e-04), indicating consistent pathway dysregulation across datasets. **F)** Contrasting pathway patterns in single-cell vs. bulk tumors: Heatmap of all Hallmark pathways comparing High-AS vs. Low-AS in TCGA (left) and single-cell (right) datasets. Color reflects enrichment results (NES for TCGA; pooled effect size for single-cell). Significant pathways ($q < 0.05$) are highlighted with a bold border.

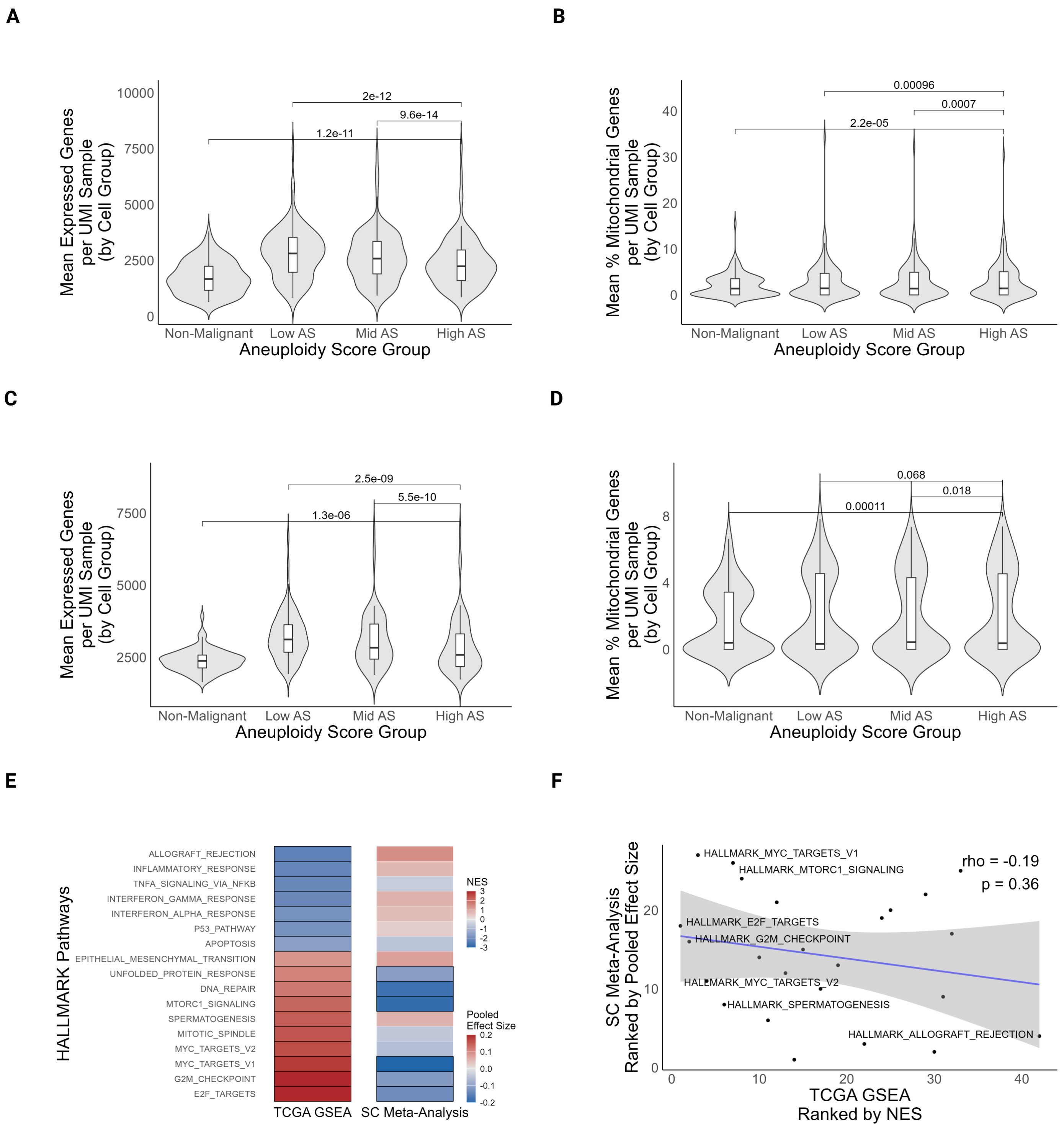
A
B
C
D
E
F
Mean Expressed Genes per UMI Sample (by Cell Group)
Mean % Mitochondrial Genes per UMI Sample (by Cell Group)
Aneuploidy Score Group
Non-Malignant
Low AS
Mid AS
High AS
2e-12
9.6e-14
1.2e-11
0.00096
0.0007
2.2e-05
2.5e-09
5.5e-10
1.3e-06
0.068
0.018
0.00011
HALLMARK Pathways
ALLOGRAFT_REJECTION
INFLAMMATORY_RESPONSE
TNFA_SIGNALING_VIA_NFKB
INTERFERON_GAMMA_RESPONSE
INTERFERON_ALPHA_RESPONSE
P53_PATHWAY
APOPTOSIS
EPITHELIAL_MESENCHYMAL_TRANSITION
UNFOLDED_PROTEIN_RESPONSE
DNA_REPAIR
MTORC1_SIGNALING
SPERMATOGENESIS
MITOTIC_SPINDLE
MYC_TARGETS_V2
MYC_TARGETS_V1
G2M_CHECKPOINT
E2F_TARGETS
TCGA GSEA
SC Meta-Analysis
NES
Pooled Effect Size
SC Meta-Analysis Ranked by Pooled Effect Size
TCGA GSEA Ranked by NES
rho = -0.19
p = 0.36
HALLMARK_MYC_TARGETS_V1
HALLMARK_MTORC1_SIGNALING
HALLMARK_E2F_TARGETS
HALLMARK_G2M_CHECKPOINT
HALLMARK_MYC_TARGETS_V2
HALLMARK_SPERMATOGENESIS
HALLMARK_ALLOGRAFT_REJECTION

**Supplementary Figure S4: Inverse relationship between single-cell and bulk-tumor gene expression is not driven by cell quality (related to Fig.2)**

**A)** Cellular gene diversity across aneuploidy groups: Violin plots comparing the mean number of expressed genes per UMI count tumor sample (n=108) across non-malignant, Low-AS, Mid-AS, and High-AS cell populations. High-AS cells exhibit a significantly lower mean number of expressed genes compared to Low-AS (p = 2e-12), Mid-AS (p = 9.6e-14), and non-malignant cell groups (p = 1.2e-11; paired two-tailed Wilcoxon signed-rank test). **B)** Mitochondrial transcript fraction across aneuploidy groups: Violin plots comparing the mean mitochondrial transcript percentage per UMI count tumor sample (n=108) across cell populations. High-AS cells exhibit a significantly higher mean mitochondrial transcript fraction compared to Low-AS (p = 0.00096), Mid-AS (p = 0.0007), and non-malignant cell groups (p=2.2e-5; paired two-tailed Wilcoxon signed-rank test). **C)** Maintenance of transcriptional breadth in high-quality cells: Violin plots comparing the mean number of expressed genes per UMI count tumor sample across cell groups after excluding lower-quality cells (cells with <1,500 genes or >10\% mitochondrial DNA, n=81). High-AS cells continue to exhibit a significantly lower mean number of expressed genes compared to Low-AS (p = 2.5e-9), Mid-AS (p = 5.5e-10), and non-malignant groups (p = 1.3e-6; paired two-tailed Wilcoxon signed-rank test). **D)** Stabilization of mitochondrial content following stringent filtering: Violin plots comparing the mean mitochondrial transcript percentage per UMI count tumor sample across cell groups in the filtered high-quality subset (n=81). No significant differences in mitochondrial fraction were observed between High-AS and Low-AS cells (paired two-tailed Wilcoxon signed-rank test, p=0.068). **E)** Differential pathway activation between TCGA and our single-cell analysis holds true within the filtered high-quality subset of cells: Selected 'Hallmark' gene sets related to proliferation, metabolism, and immunity are shown across TCGA (left) and single-cell (right) datasets, comparing High-AS versus Low-AS groups within the filtered high-quality subset (n=81). Color represents enrichment results (NES for TCGA; Polled effect size for single-cell). Significant pathways (q < 0.05) are highlighted with a bold border. **F)** Inverse correlation between single-cell and bulk tumor pathway changes within the filtered high-quality subset: Scatter plot comparing ranked pathway enrichments between single-cell and TCGA datasets within the filtered high-quality subset (n=81). The x-axis ranks TCGA 'Hallmark' pathways by NES; the y-axis ranks the same pathways by single-cell pooled effect sizes. The opposite expression trends were observed, albeit not significantly (Spearman's rho = -0.19, p=0.36).

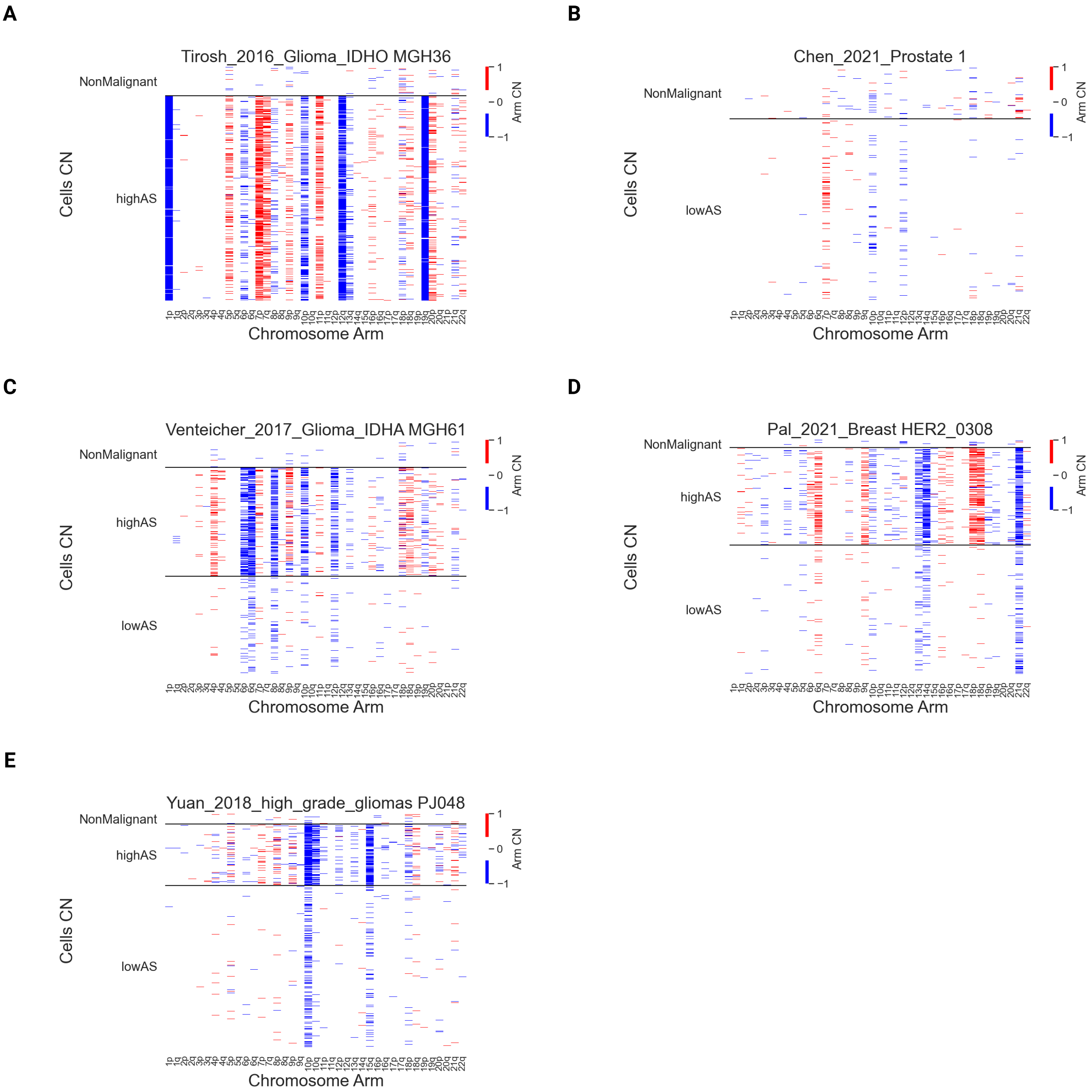
A
Tirosh_2016_Glioma_IDHO MGH36
NonMalignant
highAS
Cells CN
Chromosome Arm
Arm CN
1
0
−1
B
Chen_2021_Prostate 1
NonMalignant
lowAS
Cells CN
Chromosome Arm
Arm CN
C
Venteicher_2017_Glioma_IDHA MGH61
NonMalignant
highAS
lowAS
Cells CN
Chromosome Arm
Arm CN
D
Pal_2021_Breast HER2_0308
NonMalignant
highAS
lowAS
Cells CN
Chromosome Arm
Arm CN
E
Yuan_2018_high_grade_gliomas PJ048
NonMalignant
highAS
lowAS
Cells CN
Chromosome Arm
Arm CN

**Supplementary Figure S5: Inferred arm-level aneuploidy profiles using stringent aneuploidy scoring parameters (related to Fig. 2)**

**A-E)** Arm-level copy number profiles across diverse tumor contexts: Heatmaps showing chromosomal arm-level copy number (CN) alterations in representative tumors, including (A) highly aneuploid, (B) near-diploid, and (C-E) heterogeneous tumors containing a mixture of Low-AS and High-AS malignant cells. Each column represents a chromosomal arm; each row represents a single cell. Cells are grouped by their classification as non-malignant or malignant (Low-AS, Mid-AS, or High-AS) based on the refined, high-stringency scoring parameters. Color indicates inferred arm-level CN values (red = gain; blue = loss). Tumors are ordered in the same order that they appear in **Supplementary Fig. S2**, the difference between the plots is in the parameters used for aneuploidy calling (see **Methods**).

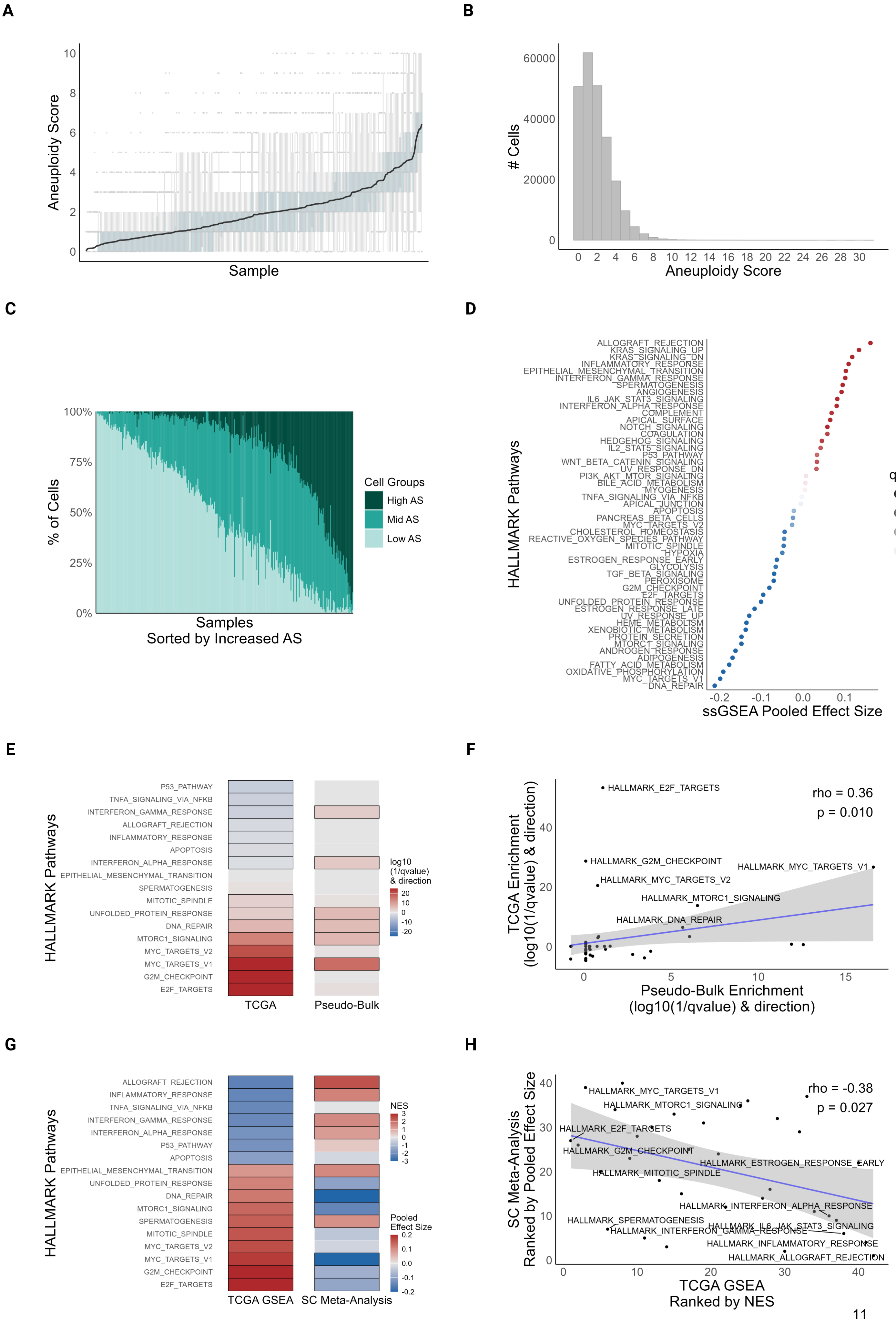
A
Aneuploidy Score
Sample
B
### Cells
Aneuploidy Score
C
% of Cells
Samples
Sorted by Increased AS
Cell Groups
High AS
Mid AS
Low AS
D
HALLMARK Pathways
ssGSEA Pooled Effect Size
q-value
ALLOGRAFT_REJECTION
KRAS_SIGNALING_UP
KRAS_SIGNALING_DN
INFLAMMATORY_RESPONSE
EPITHELIAL_MESENCHYMAL_TRANSITION
INTERFERON_GAMMA_RESPONSE
SPERMATOGENESIS
ANGIOGENESIS
IL6_JAK_STAT3_SIGNALING
INTERFERON_ALPHA_RESPONSE
COMPLEMENT
APICAL_SURFACE
NOTCH_SIGNALING
COAGULATION
HEDGEHOG_SIGNALING
IL2_STAT5_SIGNALING
P53_PATHWAY
WNT_BETA_CATENIN_SIGNALING
UV_RESPONSE_DN
PI3K_AKT_MTOR_SIGNALING
BILE_ACID_METABOLISM
MYOGENESIS
TNFA_SIGNALING_VIA_NFKB
APICAL_JUNCTION
APOPTOSIS
PANCREAS_BETA_CELLS
MYC_TARGETS_V2
CHOLESTEROL_HOMEOSTASIS
REACTIVE_OXYGEN_SPECIES_PATHWAY
MITOTIC_SPINDLE
HYPOXIA
ESTROGEN_RESPONSE_EARLY
GLYCOLYSIS
TGF_BETA_SIGNALING
PEROXISOME
G2M_CHECKPOINT
E2F_TARGETS
UNFOLDED_PROTEIN_RESPONSE
ESTROGEN_RESPONSE_LATE
UV_RESPONSE_UP
HEME_METABOLISM
XENOBIOTIC_METABOLISM
PROTEIN_SECRETION
MTORC1_SIGNALING
ANDROGEN_RESPONSE
ADIPOGENESIS
FATTY_ACID_METABOLISM
OXIDATIVE_PHOSPHORYLATION
MYC_TARGETS_V1
DNA_REPAIR
E
HALLMARK Pathways
P53_PATHWAY
TNFA_SIGNALING_VIA_NFKB
INTERFERON_GAMMA_RESPONSE
ALLOGRAFT_REJECTION
INFLAMMATORY_RESPONSE
APOPTOSIS
INTERFERON_ALPHA_RESPONSE
EPITHELIAL_MESENCHYMAL_TRANSITION
SPERMATOGENESIS
MITOTIC_SPINDLE
UNFOLDED_PROTEIN_RESPONSE
DNA_REPAIR
MTORC1_SIGNALING
MYC_TARGETS_V2
MYC_TARGETS_V1
G2M_CHECKPOINT
E2F_TARGETS
TCGA
Pseudo-Bulk
log10 (1/qvalue) & direction
F
TCGA Enrichment (log10(1/qvalue) & direction)
Pseudo-Bulk Enrichment (log10(1/qvalue) & direction)
rho = 0.36
p = 0.010
HALLMARK_E2F_TARGETS
HALLMARK_G2M_CHECKPOINT
HALLMARK_MYC_TARGETS_V1
HALLMARK_MYC_TARGETS_V2
HALLMARK_MTORC1_SIGNALING
HALLMARK_DNA_REPAIR
G
HALLMARK Pathways
ALLOGRAFT_REJECTION
INFLAMMATORY_RESPONSE
TNFA_SIGNALING_VIA_NFKB
INTERFERON_GAMMA_RESPONSE
INTERFERON_ALPHA_RESPONSE
P53_PATHWAY
APOPTOSIS
EPITHELIAL_MESENCHYMAL_TRANSITION
UNFOLDED_PROTEIN_RESPONSE
DNA_REPAIR
MTORC1_SIGNALING
SPERMATOGENESIS
MITOTIC_SPINDLE
MYC_TARGETS_V2
MYC_TARGETS_V1
G2M_CHECKPOINT
E2F_TARGETS
TCGA GSEA
SC Meta-Analysis
NES
Pooled Effect Size
H
SC Meta-Analysis Ranked by Pooled Effect Size
TCGA GSEA Ranked by NES
rho = -0.38
p = 0.027
HALLMARK_MYC_TARGETS_V1
HALLMARK_MTORC1_SIGNALING
HALLMARK_E2F_TARGETS
HALLMARK_G2M_CHECKPOINT
HALLMARK_ESTROGEN_RESPONSE_EARLY
HALLMARK_MITOTIC_SPINDLE
HALLMARK_INTERFERON_ALPHA_RESPONSE
HALLMARK_SPERMATOGENESIS
HALLMARK_IL6_JAK_STAT3_SIGNALING
HALLMARK_INTERFERON_GAMMA_RESPONSE
HALLMARK_INFLAMMATORY_RESPONSE
HALLMARK_ALLOGRAFT_REJECTION

**Supplementary Figure S6: Inverse relationship between single-cell and bulk-tumor gene expression is not affected by aneuploidy scoring parameters**

**A)** AS Distribution across tumor samples: Bar plot displaying tumor-level AS for 279 samples with background box plots illustrating the distribution of constituent malignant single-cell scores. **B)** Distribution of malignant cells by AS: Histogram representing the frequency of individual malignant cells (n = 234,744) across the AS spectrum. **C)** Stratification of cells into AS groups: Bar plot illustrating the relative distribution of cells across Low (AS ≤ 1, light green), Mid (1 < AS < 4, medium green), and High (AS ≥ 4, dark green) AS groups across 279 tumors. **D)** 'Hallmark' gene set enrichment analysis in single cells: Dot plot showing pooled effect sizes for 'Hallmark' gene sets comparing High-AS versus Low-AS cells (n = 104 tumors). Dot color indicates direction of enrichment (red = upregulated; blue = downregulated in High-AS), and transparency reflects statistical significance (q-value). **E)** 'Hallmark' gene set enrichment analysis in bulk tumors: Heatmap comparing High-AS versus Low-AS tumors in both pseudo-bulk and TCGA datasets. Black boxes denote significantly enriched Hallmark pathways related to proliferation, metabolism, and immunity (q < 0.05). **F)** Concordant pathway enrichment between bulk datasets: Scatter plot comparing enrichment scores between pseudo-bulk and TCGA tumors. A positive correlation is observed (Spearman's rho = 0.36, p = 0.01). **G)** Contrasting pathway patterns in single-cell vs. bulk tumors: Heatmap comparing selected Hallmark gene sets across TCGA (left) and single-cell (right) datasets. Color reflects enrichment results (NES for TCGA; Pooled effect size for single-cell); significant pathways (q < 0.05) are highlighted with a bold border. **H)** Inverse correlation between single-cell and bulk-tumor pathway changes: Scatter plot comparing ranked pathway enrichments between TCGA (NES) and single-cell (pooled effect size) datasets. A significant negative correlation is observed (Spearman's rho = -0.38, p = 0.027), confirming the robustness of the inverse transcriptional relationship.

A

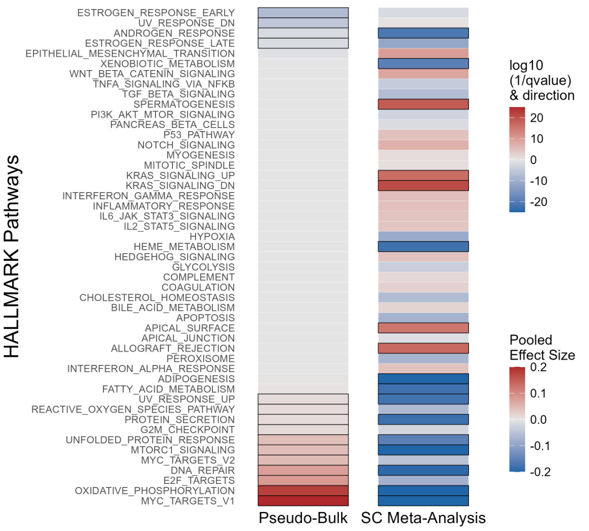
HALLMARK Pathways
ESTROGEN_RESPONSE_EARLY
UV_RESPONSE_DN
ANDROGEN_RESPONSE
ESTROGEN_RESPONSE_LATE
EPITHELIAL_MESENCHYMAL_TRANSITION
XENOBIOTIC_METABOLISM
WNT_BETA_CATENIN_SIGNALING
TNFA_SIGNALING_VIA_NFKB
TGF_BETA_SIGNALING
SPERMATOGENESIS
PI3K_AKT_MTOR_SIGNALING
PANCREAS_BETA_CELLS
P53_PATHWAY
NOTCH_SIGNALING
MYOGENESIS
MITOTIC_SPINDLE
KRAS_SIGNALING_UP
KRAS_SIGNALING_DN
INTERFERON_GAMMA_RESPONSE
INFLAMMATORY_RESPONSE
IL6_JAK_STAT3_SIGNALING
IL2_STAT5_SIGNALING
HYPOXIA
HEME_METABOLISM
HEDGEHOG_SIGNALING
GLYCOLYSIS
COMPLEMENT
COAGULATION
CHOLESTEROL_HOMEOSTASIS
BILE_ACID_METABOLISM
APOPTOSIS
APICAL_SURFACE
APICAL_JUNCTION
ALLOGRAFT_REJECTION
PEROXISOME
INTERFERON_ALPHA_RESPONSE
ADIPOGENESIS
FATTY_ACID_METABOLISM
UV_RESPONSE_UP
REACTIVE_OXYGEN_SPECIES_PATHWAY
PROTEIN_SECRETION
G2M_CHECKPOINT
UNFOLDED_PROTEIN_RESPONSE
MTORC1_SIGNALING
MYC_TARGETS_V2
DNA_REPAIR
E2F_TARGETS
OXIDATIVE_PHOSPHORYLATION
MYC_TARGETS_V1
Pseudo-Bulk
SC Meta-Analysis
log10 (1/qvalue) & direction
20
10
0
-10
-20
Pooled Effect Size
0.2
0.1
0.0
-0.1
-0.2

B

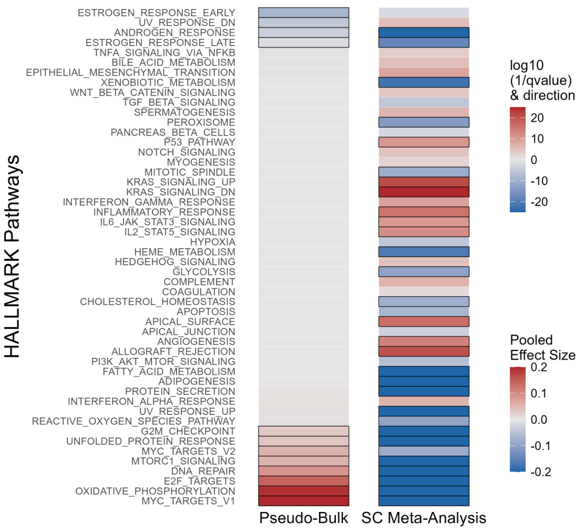
HALLMARK Pathways
ESTROGEN_RESPONSE_EARLY
UV_RESPONSE_DN
ANDROGEN_RESPONSE
ESTROGEN_RESPONSE_LATE
TNFA_SIGNALING_VIA_NFKB
BILE_ACID_METABOLISM
EPITHELIAL_MESENCHYMAL_TRANSITION
XENOBIOTIC_METABOLISM
WNT_BETA_CATENIN_SIGNALING
TGF_BETA_SIGNALING
SPERMATOGENESIS
PEROXISOME
PANCREAS_BETA_CELLS
P53_PATHWAY
NOTCH_SIGNALING
MYOGENESIS
MITOTIC_SPINDLE
KRAS_SIGNALING_UP
KRAS_SIGNALING_DN
INTERFERON_GAMMA_RESPONSE
INFLAMMATORY_RESPONSE
IL6_JAK_STAT3_SIGNALING
IL2_STAT5_SIGNALING
HYPOXIA
HEME_METABOLISM
HEDGEHOG_SIGNALING
GLYCOLYSIS
COMPLEMENT
COAGULATION
CHOLESTEROL_HOMEOSTASIS
APOPTOSIS
APICAL_SURFACE
APICAL_JUNCTION
ANGIOGENESIS
ALLOGRAFT_REJECTION
PI3K_AKT_MTOR_SIGNALING
FATTY_ACID_METABOLISM
ADIPOGENESIS
PROTEIN_SECRETION
INTERFERON_ALPHA_RESPONSE
UV_RESPONSE_UP
REACTIVE_OXYGEN_SPECIES_PATHWAY
G2M_CHECKPOINT
UNFOLDED_PROTEIN_RESPONSE
MYC_TARGETS_V2
MTORC1_SIGNALING
DNA_REPAIR
E2F_TARGETS
OXIDATIVE_PHOSPHORYLATION
MYC_TARGETS_V1
Pseudo-Bulk
SC Meta-Analysis
log10 (1/qvalue) & direction
20
10
0
-10
-20
Pooled Effect Size
0.2
0.1
0.0
-0.1
-0.2

**Supplementary Figure S7: Inverse relationship between single-cell and bulk-tumor gene expression is not due to cell cycle differences**

**A)** Contrasting pathway patterns in G1-phase cells: Heatmap comparing Hallmark pathway enrichment between pseudo-bulk (left) and single-cell meta-analysis (right) using only cells isolated in G1 phase. Analysis was restricted to tumor samples containing at least 100 G1-phase malignant cells (n = 173). Color reflects enrichment results (enrichment direction and strength for pseudo-bulk; pooled effect size for single-cell). Significant pathways ($q < 0.05$) are highlighted with a bold border. **B)** Contrasting pathway patterns in G2/M and S-phase cells: Heatmap comparing Hallmark pathway enrichment between pseudo-bulk (left) and single-cell meta-analysis (right) using only cells isolated in G2/M and S phases. Analysis was restricted to tumor samples containing at least 100 cycling malignant cells (n = 181) Color reflects enrichment results (enrichment direction and strength for pseudo-bulk; pooled effect size for single-cell). Significant pathways ($q < 0.05$) are highlighted with a bold border.

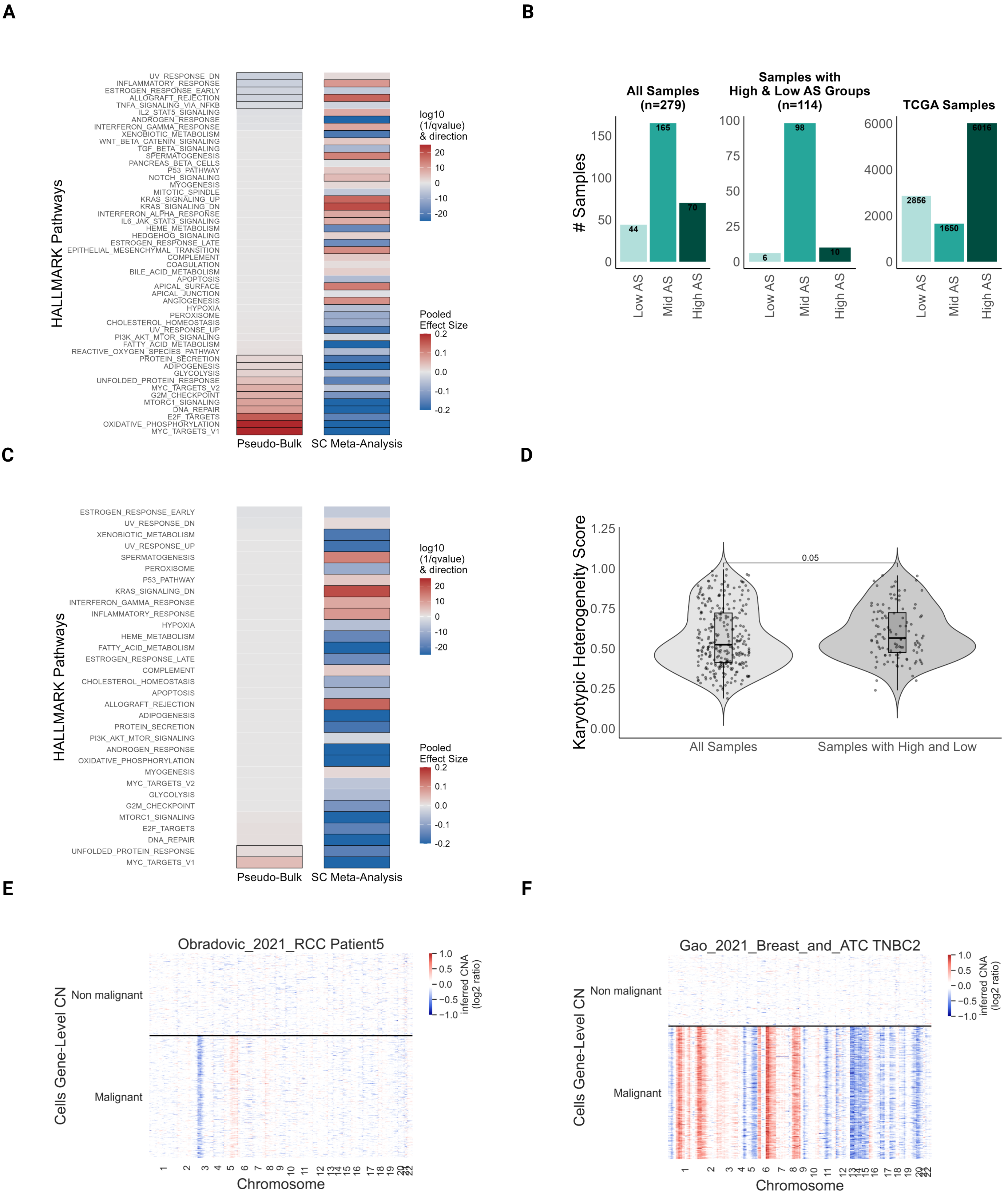
A
HALLMARK Pathways
UV_RESPONSE_DN
INFLAMMATORY_RESPONSE
ESTROGEN_RESPONSE_EARLY
ALLOGRAFT_REJECTION
TNFA_SIGNALING_VIA_NFKB
IL2_STAT5_SIGNALING
ANDROGEN_RESPONSE
INTERFERON_GAMMA_RESPONSE
XENOBIOTIC_METABOLISM
WNT_BETA_CATENIN_SIGNALING
TGF_BETA_SIGNALING
SPERMATOGENESIS
PANCREAS_BETA_CELLS
P53_PATHWAY
NOTCH_SIGNALING
MYOGENESIS
MITOTIC_SPINDLE
KRAS_SIGNALING_UP
KRAS_SIGNALING_DN
INTERFERON_ALPHA_RESPONSE
IL6_JAK_STAT3_SIGNALING
HEME_METABOLISM
HEDGEHOG_SIGNALING
ESTROGEN_RESPONSE_LATE
EPITHELIAL_MESENCHYMAL_TRANSITION
COMPLEMENT
COAGULATION
BILE_ACID_METABOLISM
APOPTOSIS
APICAL_SURFACE
APICAL_JUNCTION
ANGIOGENESIS
HYPOXIA
PEROXISOME
CHOLESTEROL_HOMEOSTASIS
UV_RESPONSE_UP
PI3K_AKT_MTOR_SIGNALING
FATTY_ACID_METABOLISM
REACTIVE_OXYGEN_SPECIES_PATHWAY
PROTEIN_SECRETION
ADIPOGENESIS
GLYCOLYSIS
UNFOLDED_PROTEIN_RESPONSE
MYC_TARGETS_V2
G2M_CHECKPOINT
MTORC1_SIGNALING
DNA_REPAIR
E2F_TARGETS
OXIDATIVE_PHOSPHORYLATION
MYC_TARGETS_V1
Pseudo-Bulk
SC Meta-Analysis
log10 (1/qvalue) & direction
20
10
0
-10
-20
Pooled Effect Size
0.2
0.1
0.0
-0.1
-0.2
B
### Samples
All Samples (n=279)
Samples with High & Low AS Groups (n=114)
TCGA Samples
44
165
70
6
98
10
2856
1650
6016
Low AS
Mid AS
High AS
C
HALLMARK Pathways
ESTROGEN_RESPONSE_EARLY
UV_RESPONSE_DN
XENOBIOTIC_METABOLISM
UV_RESPONSE_UP
SPERMATOGENESIS
PEROXISOME
P53_PATHWAY
KRAS_SIGNALING_DN
INTERFERON_GAMMA_RESPONSE
INFLAMMATORY_RESPONSE
HYPOXIA
HEME_METABOLISM
FATTY_ACID_METABOLISM
ESTROGEN_RESPONSE_LATE
COMPLEMENT
CHOLESTEROL_HOMEOSTASIS
APOPTOSIS
ALLOGRAFT_REJECTION
ADIPOGENESIS
PROTEIN_SECRETION
PI3K_AKT_MTOR_SIGNALING
ANDROGEN_RESPONSE
OXIDATIVE_PHOSPHORYLATION
MYOGENESIS
MYC_TARGETS_V2
GLYCOLYSIS
G2M_CHECKPOINT
MTORC1_SIGNALING
E2F_TARGETS
DNA_REPAIR
UNFOLDED_PROTEIN_RESPONSE
MYC_TARGETS_V1
Pseudo-Bulk
SC Meta-Analysis
log10 (1/qvalue) & direction
Pooled Effect Size
D
Karyotypic Heterogeneity Score
0.05
All Samples
Samples with High and Low
E
Obradovic_2021_RCC Patient5
Cells Gene-Level CN
Non malignant
Malignant
inferred CNA (log2 ratio)
Chromosome
F
Gao_2021_Breast_and_ATC TNBC2
Cells Gene-Level CN
Non malignant
Malignant
inferred CNA (log2 ratio)
Chromosome

**Supplementary Figure S8: Karyotypic heterogeneity underlies the inverse relationship between single-cell and bulk-cell population data (related to Fig.3)**

**A)** Pathway enrichment analysis using only malignant cells: Heatmap of 'Hallmark' gene set enrichments, comparing High-AS vs. Low-AS, based on pseudo-bulk analysis of malignant cells (left) and single-cell data (right). Color represents enrichment results (enrichment direction and strength for pseudo-bulk; pooled effect size for single-cell). Significant pathways ($q < 0.05$) are highlighted with a bold border. **B)** AS distribution across cohorts: Bar plots showing the number of tumors classified into Low (light green), Mid (medium green), and High (dark green) AS groups across three datasets: all tumors analyzed (left, n = 279), tumors containing both High-AS and Low-AS cells (middle, n = 114), and TCGA tumors (right, n = 10,522). **C)** Pathway enrichment analysis restricted to tumors with both High-AS and Low-AS cells: Heatmap comparing High-AS vs. Low-AS groups for Hallmark pathways in pseudo-bulk restricted to tumors containing both High and Low AS (left) and single-cell (right) analyses. Color indicates enrichment results (enrichment direction and strength for pseudo-bulk; pooled effect size for single-cell). Significant pathways ($q < 0.05$) are highlighted with a bold border. **D)** Comparison of karyotypic heterogeneity in tumors with both High-AS and Low-AS cells: Violin plots comparing tumor-level karyotypic heterogeneity between all tumors (n = 279) and the subset of tumors containing both High-AS and Low-AS cells (n = 114). Tumors with both High-AS and Low-AS cells exhibit significantly higher karyotypic heterogeneity (two-tailed Wilcoxon rank-sum test, $p = 0.0025$). **E)** Copy number profiles of a representative karyotypically heterogeneous tumor: Heatmaps showing gene-level copy number (CN) alterations in a representative karyotypically heterogeneous tumor. Each column represents a gene; each row represents a single tumor cell. Color indicates inferred CNA (log2 ratio) values (blue = loss; red = gain). **F)** Copy number profiles of representative karyotypically homogeneous tumor: Heatmaps as in (E).

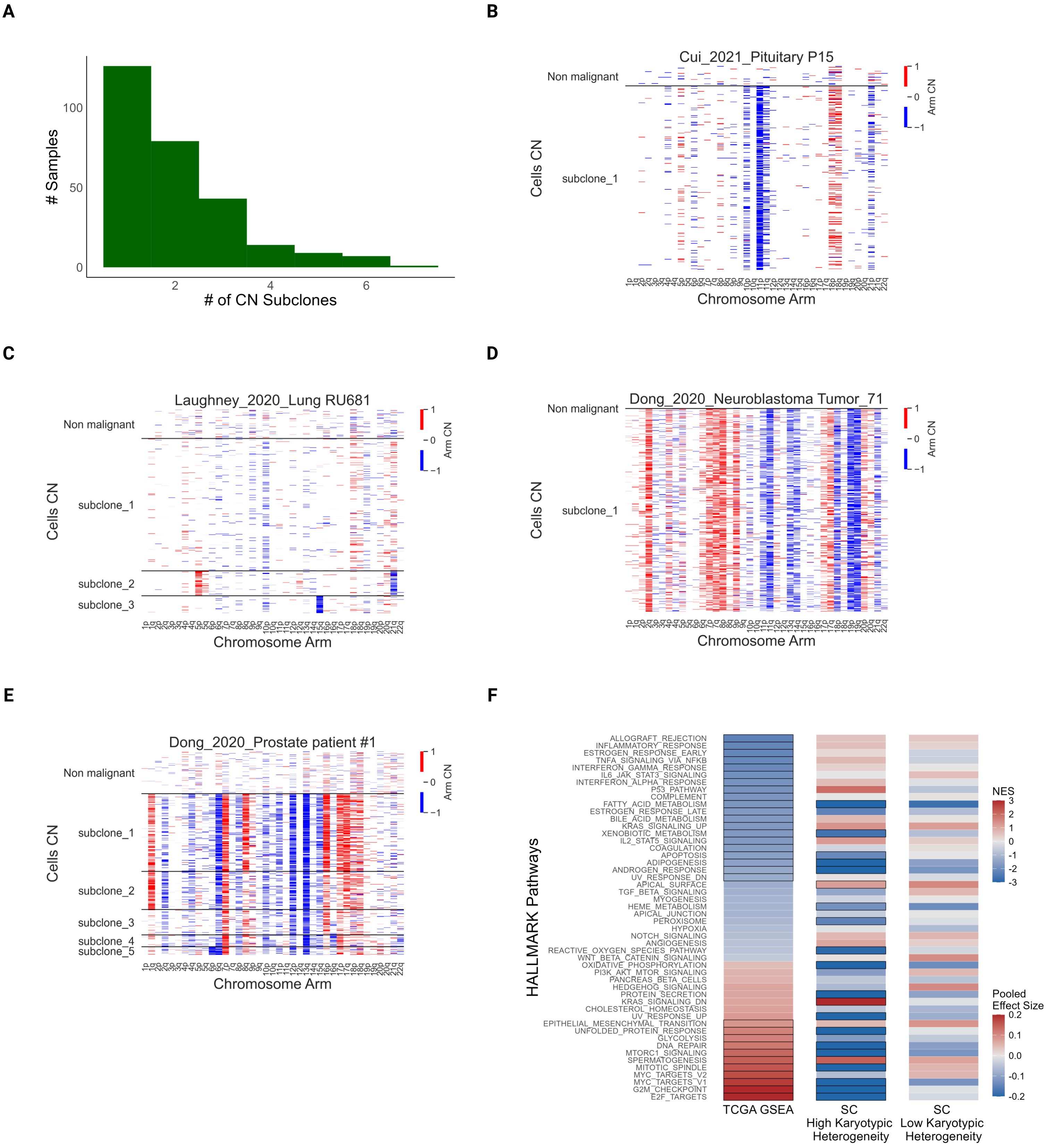

A
B
C
D
E
F
### Samples
### of CN Subclones
Cui_2021_Pituitary P15
Non malignant
subclone_1
Cells CN
Chromosome Arm
Arm CN
Laughney_2020_Lung RU681
subclone_2
subclone_3
Dong_2020_Neuroblastoma Tumor_71
Dong_2020_Prostate patient #1
subclone_4
subclone_5
HALLMARK Pathways
ALLOGRAFT_REJECTION
INFLAMMATORY_RESPONSE
ESTROGEN_RESPONSE_EARLY
TNFA_SIGNALING_VIA_NFKB
INTERFERON_GAMMA_RESPONSE
IL6_JAK_STAT3_SIGNALING
INTERFERON_ALPHA_RESPONSE
P53_PATHWAY
COMPLEMENT
FATTY_ACID_METABOLISM
ESTROGEN_RESPONSE_LATE
BILE_ACID_METABOLISM
KRAS_SIGNALING_UP
XENOBIOTIC_METABOLISM
IL2_STAT5_SIGNALING
COAGULATION
APOPTOSIS
ADIPOGENESIS
ANDROGEN_RESPONSE
UV_RESPONSE_DN
APICAL_SURFACE
TGF_BETA_SIGNALING
MYOGENESIS
HEME_METABOLISM
APICAL_JUNCTION
PEROXISOME
HYPOXIA
NOTCH_SIGNALING
ANGIOGENESIS
REACTIVE_OXYGEN_SPECIES_PATHWAY
WNT_BETA_CATENIN_SIGNALING
OXIDATIVE_PHOSPHORYLATION
PI3K_AKT_MTOR_SIGNALING
PANCREAS_BETA_CELLS
HEDGEHOG_SIGNALING
PROTEIN_SECRETION
KRAS_SIGNALING_DN
CHOLESTEROL_HOMEOSTASIS
UV_RESPONSE_UP
EPITHELIAL_MESENCHYMAL_TRANSITION
UNFOLDED_PROTEIN_RESPONSE
GLYCOLYSIS
DNA_REPAIR
MTORC1_SIGNALING
SPERMATOGENESIS
MITOTIC_SPINDLE
MYC_TARGETS_V2
MYC_TARGETS_V1
G2M_CHECKPOINT
E2F_TARGETS
TCGA GSEA
SC High Karyotypic Heterogeneity
SC Low Karyotypic Heterogeneity
NES
Pooled Effect Size

**Supplementary Figure S9: Genetic subclone diversity and gene sets stratification by karyotypic heterogeneity (related to Fig.3)**

**A)** Distribution of genetic subclones per tumor: Histogram showing the number of genetic subclones per tumor, identified using Louvain clustering of copy number profiles. The x-axis represents the number of subclones per tumor; the y-axis indicates the frequency of tumors with each subclone count. **B-E)** Subclonal arm-level copy number profiles: Heatmaps showing chromosomal arm-level copy number (CN) alterations in representative tumors, including a near-diploid tumor with (B) a single malignant subclone and (C) multiple subclones, and a highly aneuploid tumor with (D) a single subclone and (E) multiple subclones. Rows represent single cells grouped by classification as non-malignant or malignant (partitioned into subclones); columns represent chromosomal arms. Color indicates inferred arm-level CN values (red = gain; blue = loss). **F)** Pathway enrichment analysis in homogeneous vs. heterogeneous tumors: 'Hallmark' gene set enrichments, comparing High-AS vs. Low-AS tumors or cells across TCGA (left) and single-cell (SC) datasets. In SC data, results are stratified by tumor karyotypic heterogeneity: heterogeneous tumors (middle) and homogeneous tumors (right). Color represents enrichment results (NES for TCGA; pooled effect size for SC). Significant pathways ($q < 0.05$) are highlighted with a bold border.

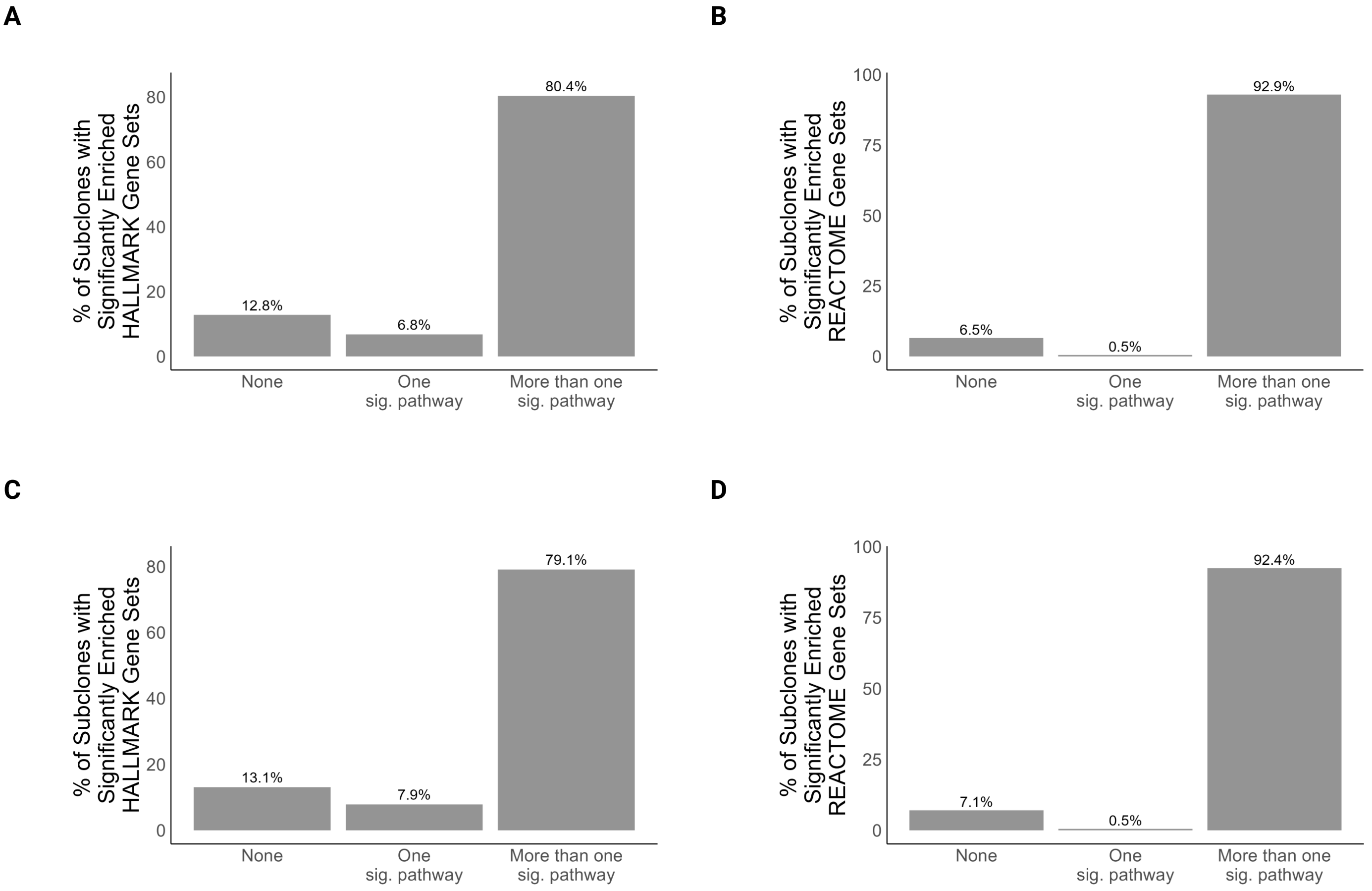
A
% of Subclones with Significantly Enriched HALLMARK Gene Sets
0
20
40
60
80
12.8%
6.8%
80.4%
None
One sig. pathway
More than one sig. pathway
B
% of Subclones with Significantly Enriched REACTOME Gene Sets
0
25
50
75
100
6.5%
0.5%
92.9%
None
One sig. pathway
More than one sig. pathway
C
% of Subclones with Significantly Enriched HALLMARK Gene Sets
0
20
40
60
80
13.1%
7.9%
79.1%
None
One sig. pathway
More than one sig. pathway
D
% of Subclones with Significantly Enriched REACTOME Gene Sets
0
25
50
75
100
7.1%
0.5%
92.4%
None
One sig. pathway
More than one sig. pathway

**Supplementary Figure S10: Gene set enrichment and depletion across genetic subclones (related to Fig.4)**

**A)** Upregulation of ‘Hallmark’ gene sets in subclones: Bar plot showing the proportion of genetic subclones (identified by Louvain clustering) in which the upregulated genes are enriched for zero, one, or more than one ‘Hallmark’ gene set relative to other subclones within the same tumor. The x-axis represents the number of significantly enriched gene sets per subclone; the y-axis shows the percentage of subclones. Approximately 80% of subclones exhibit enrichment for at least one ‘Hallmark’ gene set. **B)** Upregulation of ‘Reactome’ gene sets in subclones: Same as (A), but using ‘Reactome’ gene sets. Approximately 93% of subclones exhibit enrichment for at least one ‘Reactome’ gene set. **C)** Downregulation of ‘Hallmark’ gene sets in subclones: Bar plot showing the proportion of genetic subclones (identified by Louvain clustering) in which the downregulated genes are enriched for zero, one, or more than one ‘Hallmark’ gene set relative to other subclones within the same tumor. The x-axis represents the number of significantly enriched gene sets per subclone; the y-axis shows the percentage of subclones. Approximately 79% of subclones exhibit enrichment for at least one ‘Hallmark’ gene set. **D)** Downregulation of ‘Reactome’ gene sets in subclones: Same as (C), but using ‘Reactome’ gene sets. Approximately 92% of subclones exhibit enrichment for at least one ‘Reactome’ gene set.

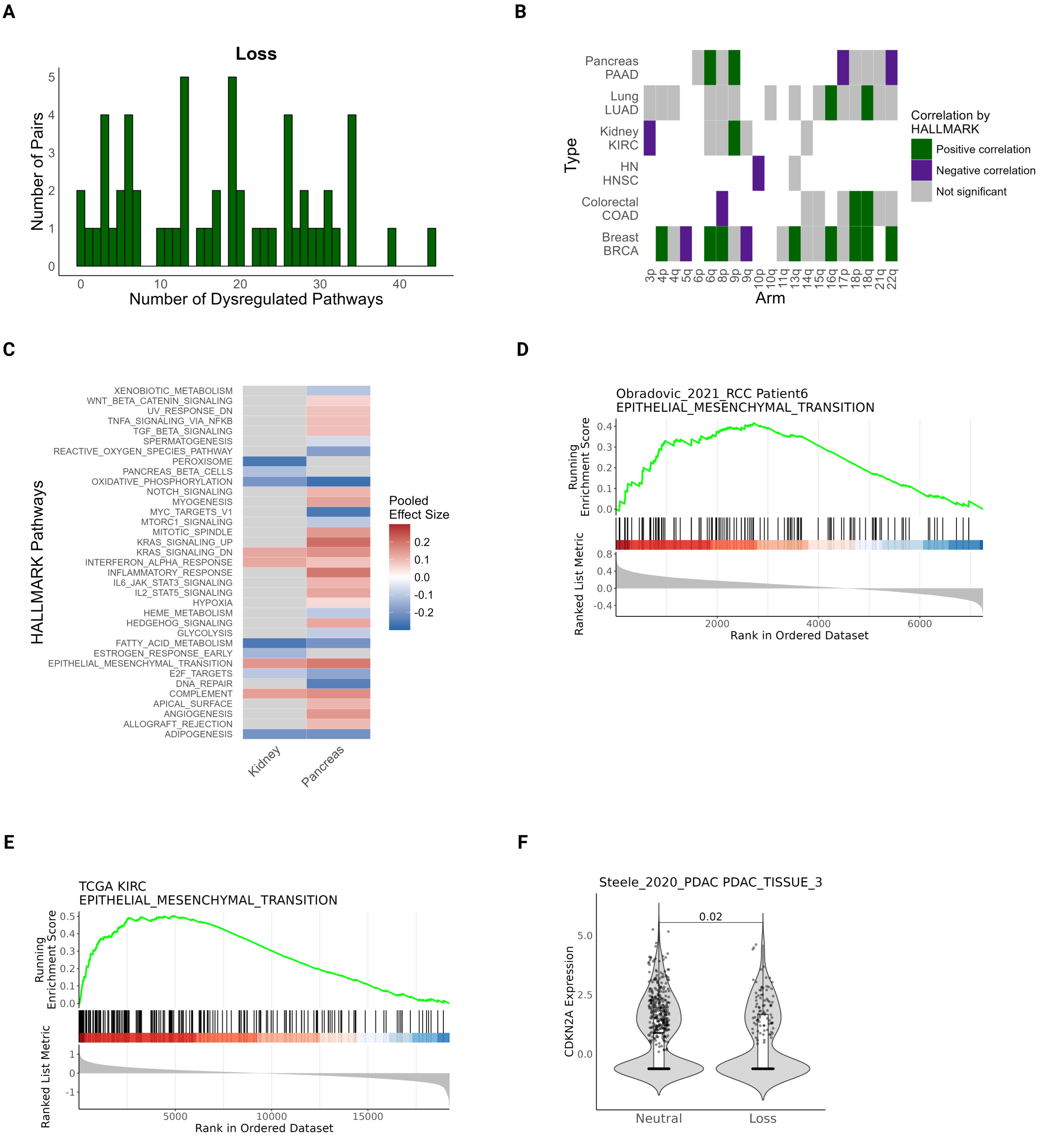
A
Loss
Number of Pairs
Number of Dysregulated Pathways
B
Pancreas PAAD
Lung LUAD
Kidney KIRC
HN HNSC
Colorectal COAD
Breast BRCA
Type
Arm
3p 4p 4q 5q 6p 6q 8p 9p 9q 10p 10q 11q 13q 14q 15q 16q 17p 18p 18q 21q 22q
Correlation by HALLMARK
Positive correlation
Negative correlation
Not significant
C
HALLMARK Pathways
XENOBIOTIC_METABOLISM
WNT_BETA_CATENIN_SIGNALING
UV_RESPONSE_DN
TNFA_SIGNALING_VIA_NFKB
TGF_BETA_SIGNALING
SPERMATOGENESIS
REACTIVE_OXYGEN_SPECIES_PATHWAY
PEROXISOME
PANCREAS_BETA_CELLS
OXIDATIVE_PHOSPHORYLATION
NOTCH_SIGNALING
MYOGENESIS
MYC_TARGETS_V1
MTORC1_SIGNALING
MITOTIC_SPINDLE
KRAS_SIGNALING_UP
KRAS_SIGNALING_DN
INTERFERON_ALPHA_RESPONSE
INFLAMMATORY_RESPONSE
IL6_JAK_STAT3_SIGNALING
IL2_STAT5_SIGNALING
HYPOXIA
HEME_METABOLISM
HEDGEHOG_SIGNALING
GLYCOLYSIS
FATTY_ACID_METABOLISM
ESTROGEN_RESPONSE_EARLY
EPITHELIAL_MESENCHYMAL_TRANSITION
E2F_TARGETS
DNA_REPAIR
COMPLEMENT
APICAL_SURFACE
ANGIOGENESIS
ALLOGRAFT_REJECTION
ADIPOGENESIS
Kidney
Pancreas
Pooled Effect Size
D
Obradovic_2021_RCC Patient6
EPITHELIAL_MESENCHYMAL_TRANSITION
Running Enrichment Score
Ranked List Metric
Rank in Ordered Dataset
E
TCGA KIRC
EPITHELIAL_MESENCHYMAL_TRANSITION
Running Enrichment Score
Ranked List Metric
Rank in Ordered Dataset
F
Steele_2020_PDAC PDAC_TISSUE_3
CDKN2A Expression
0.02
Neutral
Loss

**Supplementary Figure S11: Consistent transcriptional changes and EMT pathway activation in Chr9p-loss single-cells and tumors (related to Fig.4)**

**A)** Distribution of the number of dysregulated ‘Hallmark’ gene sets across chromosome-arm-loss – cancer-type pairs: Histogram showing the number of significantly dysregulated ‘Hallmark’ gene sets per chromosome arm–cancer-type (CA–CT) pair (for chromosome-arms with recurrent losses). The x-axis represents the number of dysregulated gene sets; the y-axis indicates the number of CA–CT pairs. **B)** Concordant transcriptional changes in TCGA and single-cell datasets: Heatmap showing correlations between transcriptional changes (‘Hallmark’ gene sets) in TCGA and single-cell datasets across 6 cancer types and 21 recurrent chromosome-arm losses (totaling 56 CA–CT pairs). Colors represent correlation direction: green, significant positive correlation, purple, significant negative correlation, gray, not significant. **C)** Gene set dysregulation associated with Chr9p loss in kidney and pancreatic single cells: Heatmap comparing ‘Hallmark’ gene set enrichment in Chr9p-loss cells vs. Chr9p-WT cells across kidney and pancreas tumors. Colors represent pooled ssGSEA effect sizes. Gray indicates non-significant results ($q \geq 0.25$). **D)** Example GSEA plot of EMT upregulation in a Chr9p-loss kidney tumor: Representative Gene Set Enrichment Analysis (GSEA) plot showing EMT gene set enrichment in Chr9p-loss cells vs. Chr9p-WT cells in a kidney tumor. **E)** EMT pathway is upregulated in TCGA kidney tumors with Chr9p loss: GSEA plot showing EMT pathway enrichment in Chr9p-loss tumors vs. Chr9p-WT tumors in TCGA KIRC (kidney) samples, consistent with the single-cell observations. **F)** *CDKN2A* expression is reduced in Chr9p-loss cells (pancreatic tumor example): Violin plots comparing *CDKN2A* expression between Chr9p-loss cells and Chr9p-WT cells in a representative pancreatic tumor. A significant reduction in expression is observed in Chr9p-loss cells (two-tailed Wilcoxon signed-rank test, $p = 0.02$).

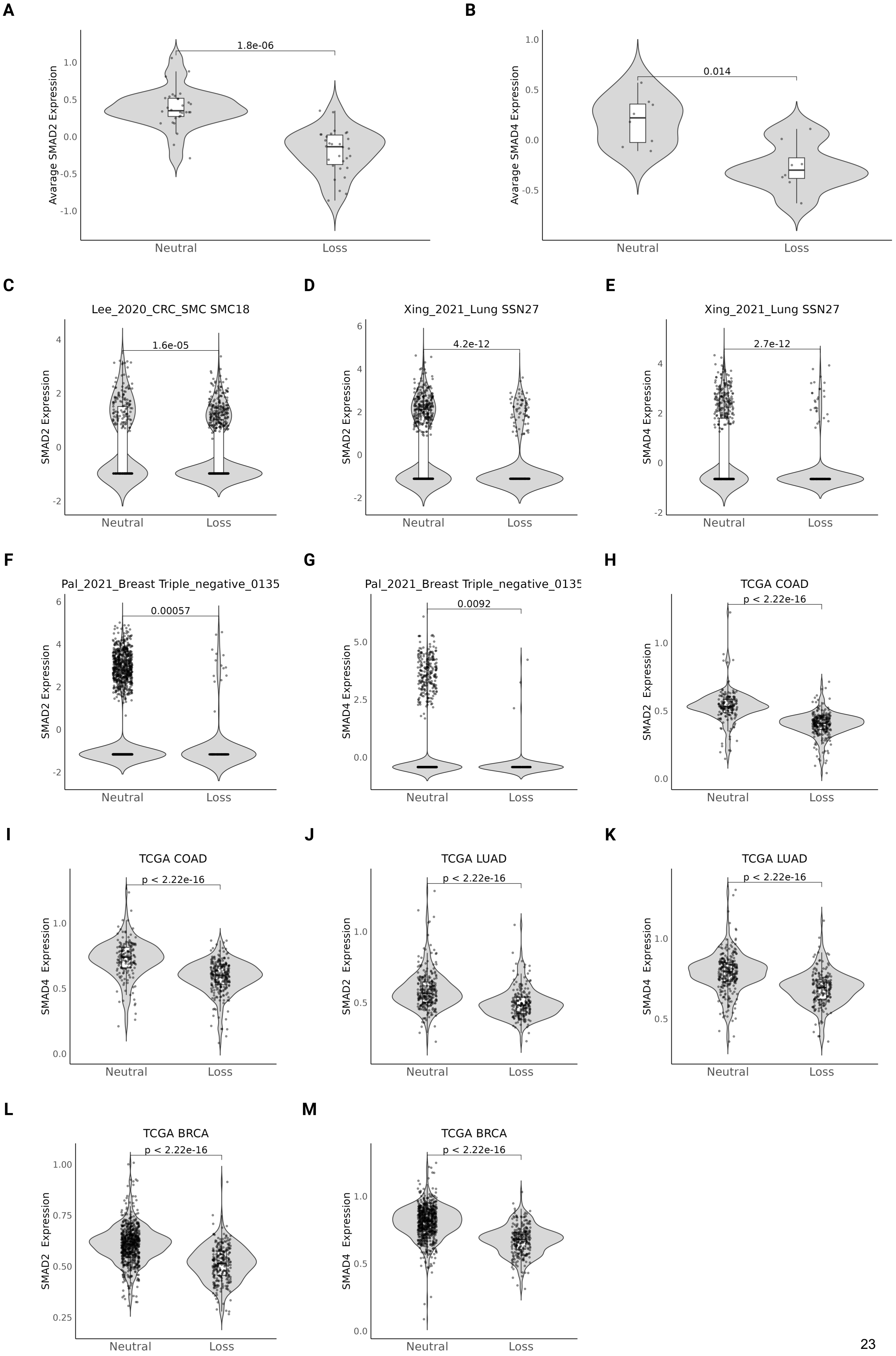
A
1.8e-06
Avarage SMAD2 Expression
Neutral
Loss
B
0.014
Avarage SMAD4 Expression
Neutral
Loss
C
Lee_2020_CRC_SMC SMC18
1.6e-05
SMAD2 Expression
D
Xing_2021_Lung SSN27
4.2e-12
SMAD2 Expression
E
Xing_2021_Lung SSN27
2.7e-12
SMAD4 Expression
F
Pal_2021_Breast Triple_negative_0135
0.00057
SMAD2 Expression
G
Pal_2021_Breast Triple_negative_0135
0.0092
SMAD4 Expression
H
TCGA COAD
p < 2.22e-16
SMAD2 Expression
I
TCGA COAD
p < 2.22e-16
SMAD4 Expression
J
TCGA LUAD
p < 2.22e-16
SMAD2 Expression
K
TCGA LUAD
p < 2.22e-16
SMAD4 Expression
L
TCGA BRCA
p < 2.22e-16
SMAD2 Expression
M
TCGA BRCA
p < 2.22e-16
SMAD4 Expression

**Supplementary Figure S12: SMAD2/4 downregulation in Chr18q-loss single-cells and tumors (related to Fig.4)**

**A)** *SMAD2* expression is reduced in Chr18q-loss cells: Violin plots comparing average *SMAD2* expression between Chr18q-loss cells and Chr18q-WT cells across 30 tumors (colorectal, lung, and breast). A significant reduction in *SMAD2* expression is observed in Chr18q-loss cells (paired two-tailed Wilcoxon signed-rank test, p = 1.8e–06). **B)** *SMAD4* expression is reduced in Chr18q-loss cells: Same as (A), but for *SMAD4* gene expression across 8 lung and breast tumors. A significant reduction is observed in Chr18q-loss cells (paired two-tailed Wilcoxon signed-rank test, p = 0.014). **C)** *SMAD2* downregulation in a representative colorectal tumor: Violin plot showing reduced *SMAD2* expression in Chr18q-loss cells vs.Chr18q-WT cells from a representative colorectal tumor (two-tailed Wilcoxon signed-rank test, p = 1.6e–05). **D)** *SMAD2* downregulation in a representative lung tumor: Same as (C), but for a representative lung tumor (two-tailed Wilcoxon signed-rank test, p = 4.2e–12). **E)** *SMAD4* downregulation in a representative lung tumor: Violin plot comparing *SMAD4* expression in Chr18q-loss cells vs. Chr18q-WT cells in a representative lung tumor (two-tailed Wilcoxon signed-rank test, p = 2.7e–12). **F)** *SMAD2* downregulation in a representative breast tumor: Same as (C), but for a representative breast tumor (two-tailed Wilcoxon signed-rank test, p = 0.00057). **G)** *SMAD4* downregulation in a representative breast tumor: Same as (E), but for a representative breast tumor (two-tailed Wilcoxon signed-rank test, p = 0.0092). **H)** *SMAD2* expression is reduced in TCGA colorectal tumors with Chr18q loss: Violin plot comparing *SMAD2* expression in Chr18q-loss vs. Chr18q-WT colorectal (COAD) tumors from the TCGA data set (two-tailed Wilcoxon rank-sum test, p < 2.2e–16). **I)** *SMAD4* expression is reduced in TCGA colorectal tumors with Chr18q loss: Same as (H), but for *SMAD4* gene (two-tailed Wilcoxon rank-sum test, p < 2.2e–16). **J)** *SMAD2* expression is reduced in TCGA lung tumors with Chr18q loss: Violin plot showing reduced *SMAD2* expression in Chr18q-loss vs. Chr18q-WT lung adenocarcinoma (LUAD) tumors from the TCGA data set (two-tailed Wilcoxon rank-sum test, p < 2.2e–16). **K)** *SMAD4* expression is reduced in TCGA lung tumors with Chr18q loss: Same as (J), but for *SMAD4* (two-tailed Wilcoxon rank-sum test, p < 2.2e–16). **L)** *SMAD2* expression is reduced in TCGA breast tumors with Chr18q loss: Violin plot comparing *SMAD2* expression in Chr18q-loss vs. Chr18q-WT breast cancer (BRCA) tumors from the TCGA data set (two-tailed Wilcoxon rank-sum test, p < 2.2e–16). **M**) *SMAD4* expression is reduced in TCGA breast tumors with Chr18q loss: Same as (M), but for *SMAD4* (two-tailed Wilcoxon rank-sum test, p < 2.2e–16).

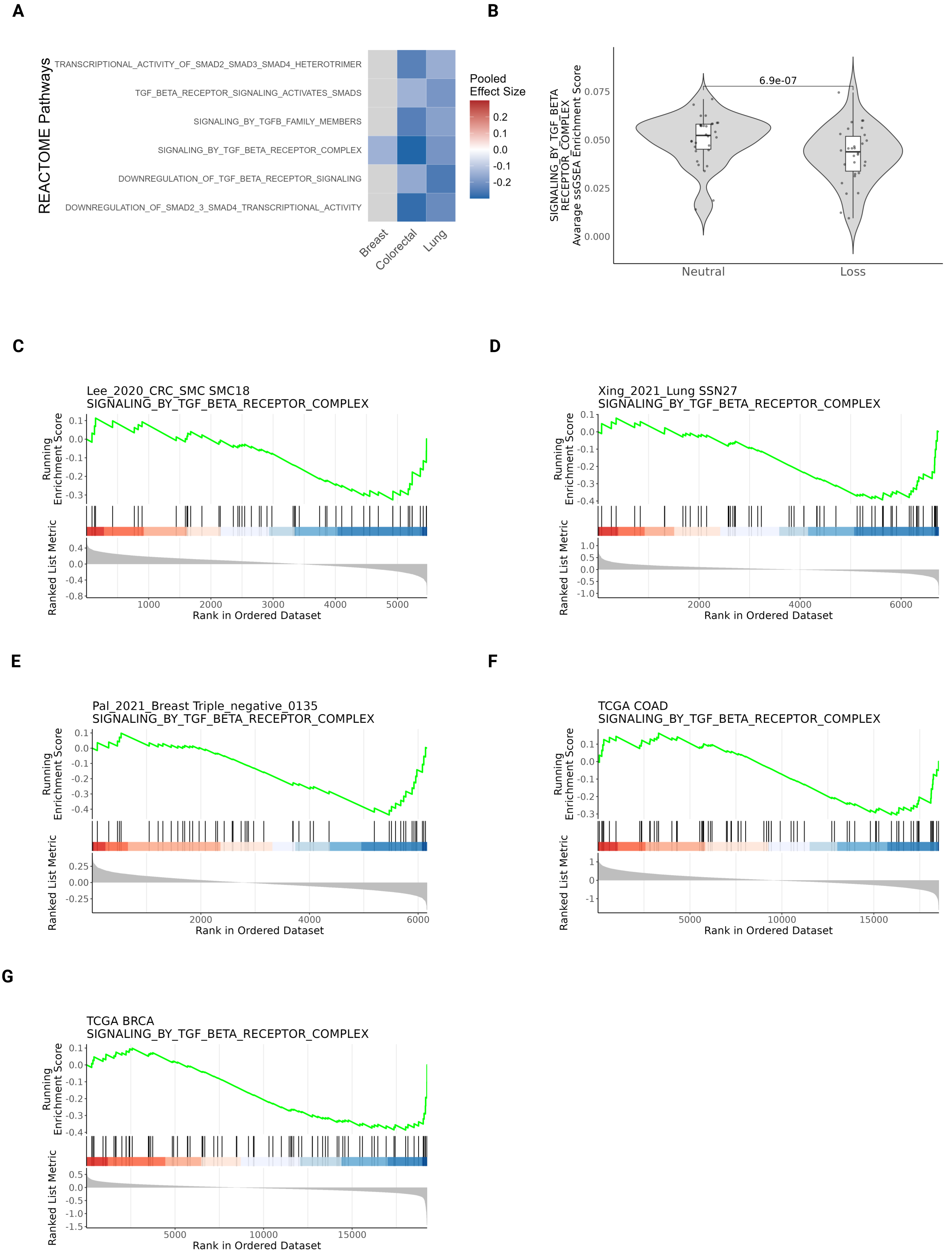
A
REACTOME Pathways
TRANSCRIPTIONAL_ACTIVITY_OF_SMAD2_SMAD3_SMAD4_HETEROTRIMER
TGF_BETA_RECEPTOR_SIGNALING_ACTIVATES_SMADS
SIGNALING_BY_TGFB_FAMILY_MEMBERS
SIGNALING_BY_TGF_BETA_RECEPTOR_COMPLEX
DOWNREGULATION_OF_TGF_BETA_RECEPTOR_SIGNALING
DOWNREGULATION_OF_SMAD2_3_SMAD4_TRANSCRIPTIONAL_ACTIVITY
Breast
Colorectal
Lung
Pooled Effect Size
0.2
0.1
0.0
-0.1
-0.2
B
SIGNALING_BY_TGF_BETA RECEPTOR_COMPLEX Avarage ssGSEA Enrichment Score
6.9e-07
0.075
0.050
0.025
0.000
Neutral
Loss
C
Lee_2020_CRC_SMC SMC18
SIGNALING_BY_TGF_BETA_RECEPTOR_COMPLEX
Running Enrichment Score
Ranked List Metric
Rank in Ordered Dataset
D
Xing_2021_Lung SSN27
SIGNALING_BY_TGF_BETA_RECEPTOR_COMPLEX
Running Enrichment Score
Ranked List Metric
Rank in Ordered Dataset
E
Pal_2021_Breast Triple_negative_0135
SIGNALING_BY_TGF_BETA_RECEPTOR_COMPLEX
Running Enrichment Score
Ranked List Metric
Rank in Ordered Dataset
F
TCGA COAD
SIGNALING_BY_TGF_BETA_RECEPTOR_COMPLEX
Running Enrichment Score
Ranked List Metric
Rank in Ordered Dataset
G
TCGA BRCA
SIGNALING_BY_TGF_BETA_RECEPTOR_COMPLEX
Running Enrichment Score
Ranked List Metric
Rank in Ordered Dataset

**Supplementary Figure S13: TGF-β pathway downregulation in Chr18q-loss cells and tumors (related to Fig.4)**

**A)** Gene set dysregulation in Chr18q-loss cells across tumor types: Heatmap showing enrichment scores for 'Reactome' pathways related to SMAD2/3/4 and TGF-β signaling in Chr18q-loss cells vs. Chr18q-WT cells across colorectal, lung, and breast tumors. Color represents pooled ssGSEA effect size. Gray indicates non-significant enrichment ($q \geq 0.25$). **B)** TGF-β signaling is reduced in Chr18q-loss cells: Violin plots comparing enrichment scores (via ssGSEA) of the 'Signaling by TGF Beta Receptor Complex' gene set between Chr18q-loss and Chr18q-WT cells across 30 tumors (colorectal, lung, and breast). A significant decrease is observed in Chr18q-loss cells gene set (paired two-tailed Wilcoxon signed-rank test, p = 6.9e–07). **C)** TGF-β signaling downregulation in a representative colorectal tumor: GSEA plot showing significant ($q<0.25$) downregulation of the 'Signaling by TGF Beta Receptor Complex' gene set in Chr18q-loss vs. Chr18q-WT cells in a representative colorectal tumor. **D)** TGF-β signaling downregulation in a representative lung tumor: Same as (C), but for a representative lung tumor. **E)** TGF-β signaling downregulation in a representative breast tumor: Same as (C), but for a representative breast tumor. **F)** Reduced TGF-β signaling in TCGA colorectal tumors with Chr18q loss: GSEA plot showing significant ($q<0.25$) downregulation of the 'Signaling by TGF Beta Receptor Complex' gene set in Chr18q-loss vs. Chr18q-WT colorectal (COAD) tumors in the TCGA data set. **G)** Reduced TGF-β signaling in TCGA breast tumors with Chr18q loss: Same as (F), but for TCGA BRCA (breast cancer) tumors.

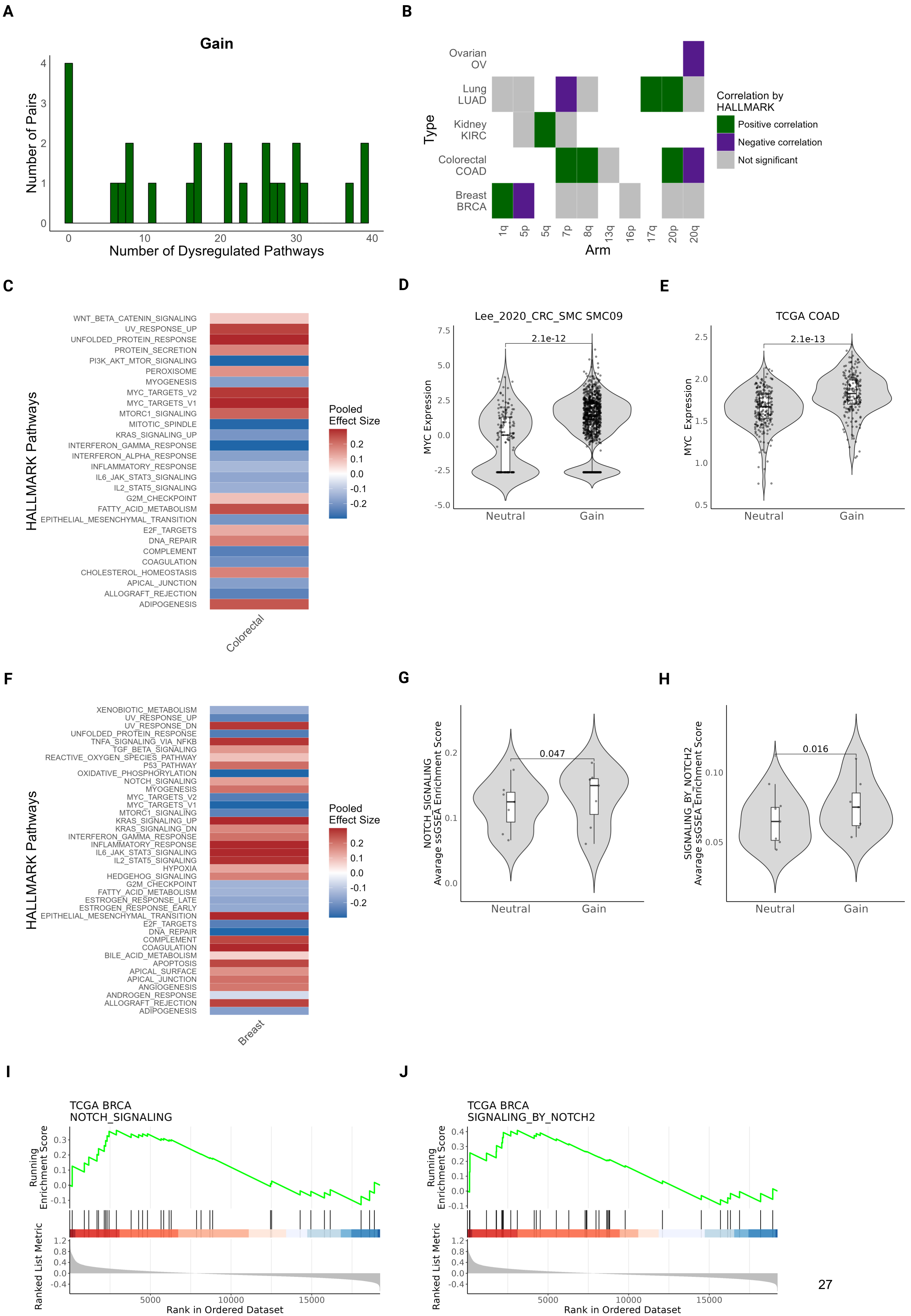
A
Gain
Number of Pairs
Number of Dysregulated Pathways
B
Ovarian OV
Lung LUAD
Kidney KIRC
Colorectal COAD
Breast BRCA
Type
1q
5p
5q
7p
8q
13q
16p
17q
20p
20q
Arm
Correlation by HALLMARK
Positive correlation
Negative correlation
Not significant
C
HALLMARK Pathways
WNT_BETA_CATENIN_SIGNALING
UV_RESPONSE_UP
UNFOLDED_PROTEIN_RESPONSE
PROTEIN_SECRETION
PI3K_AKT_MTOR_SIGNALING
PEROXISOME
MYOGENESIS
MYC_TARGETS_V2
MYC_TARGETS_V1
MTORC1_SIGNALING
MITOTIC_SPINDLE
KRAS_SIGNALING_UP
INTERFERON_GAMMA_RESPONSE
INTERFERON_ALPHA_RESPONSE
INFLAMMATORY_RESPONSE
IL6_JAK_STAT3_SIGNALING
IL2_STAT5_SIGNALING
G2M_CHECKPOINT
FATTY_ACID_METABOLISM
EPITHELIAL_MESENCHYMAL_TRANSITION
E2F_TARGETS
DNA_REPAIR
COMPLEMENT
COAGULATION
CHOLESTEROL_HOMEOSTASIS
APICAL_JUNCTION
ALLOGRAFT_REJECTION
ADIPOGENESIS
Colorectal
Pooled Effect Size
D
Lee_2020_CRC_SMC SMC09
MYC Expression
2.1e-12
Neutral
Gain
E
TCGA COAD
MYC Expression
2.1e-13
Neutral
Gain
F
HALLMARK Pathways
XENOBIOTIC_METABOLISM
UV_RESPONSE_UP
UV_RESPONSE_DN
UNFOLDED_PROTEIN_RESPONSE
TNFA_SIGNALING_VIA_NFKB
TGF_BETA_SIGNALING
REACTIVE_OXYGEN_SPECIES_PATHWAY
P53_PATHWAY
OXIDATIVE_PHOSPHORYLATION
NOTCH_SIGNALING
MYOGENESIS
MYC_TARGETS_V2
MYC_TARGETS_V1
MTORC1_SIGNALING
KRAS_SIGNALING_UP
KRAS_SIGNALING_DN
INTERFERON_GAMMA_RESPONSE
INFLAMMATORY_RESPONSE
IL6_JAK_STAT3_SIGNALING
IL2_STAT5_SIGNALING
HYPOXIA
HEDGEHOG_SIGNALING
G2M_CHECKPOINT
FATTY_ACID_METABOLISM
ESTROGEN_RESPONSE_LATE
ESTROGEN_RESPONSE_EARLY
EPITHELIAL_MESENCHYMAL_TRANSITION
E2F_TARGETS
DNA_REPAIR
COMPLEMENT
COAGULATION
BILE_ACID_METABOLISM
APOPTOSIS
APICAL_SURFACE
APICAL_JUNCTION
ANGIOGENESIS
ANDROGEN_RESPONSE
ALLOGRAFT_REJECTION
ADIPOGENESIS
Breast
Pooled Effect Size
G
NOTCH_SIGNALING Avarage ssGSEA Enrichment Score
0.047
Neutral
Gain
H
SIGNALING_BY_NOTCH2 Avarage ssGSEA Enrichment Score
0.016
Neutral
Gain
I
TCGA BRCA
NOTCH_SIGNALING
Running Enrichment Score
Ranked List Metric
Rank in Ordered Dataset
J
TCGA BRCA
SIGNALING_BY_NOTCH2
Running Enrichment Score
Ranked List Metric
Rank in Ordered Dataset

**Supplementary Figure S14: Consistent transcriptional changes, Chr8q-gain-associated MYC pathway activation, and Chr1q-gain-associated Notch signaling activation, in cells and tumors (related to Fig.4)**

**A)** Distribution of the number of dysregulated 'Hallmark' gene sets across chromosome-arm-gain–cancer-type pairs: Histogram showing the number of significantly dysregulated 'Hallmark' gene sets per chromosome arm–cancer-type (CA–CT) pair (for chromosome-arms with recurrent gains). The x-axis represents the number of dysregulated gene sets; the y-axis indicates the number of CA–CT pairs. **B)** Concordant transcriptional changes in TCGA and single-cell datasets: Heatmap showing correlations between transcriptional changes ('Hallmark gene sets) in TCGA and single-cell datasets across 5 cancer types and 10 recurrent chromosome-arm gains (totaling 23 CA–CT pairs). Colors represent correlation direction: green, significant positive correlation, purple, significant negative correlation, gray, not significant. **C)** Gene set dysregulation associated with Chr8q gain in colorectal single cells: Heatmap showing significantly enriched 'Hallmark' gene sets in Chr8q-gain vs. Chr8q-WT colorectal tumor cells. Colors represent pooled ssGSEA effect sizes. **D)** *MYC* expression is increased in Chr8q-gain colorectal cells: Violin plot comparing *MYC* expression in Chr8q-gain vs. Chr8q-WT cells in a representative colorectal tumor. *MYC* expression is significantly upregulated in Chr8q-gain cells (two-tailed Wilcoxon signed-rank test, p = 2.1e–12). **E)** *MYC* expression is increased in TCGA colorectal tumors with Chr8q gain: Violin plot showing significantly increased *MYC* expression in Chr8q-gain vs. Chr8q-WT colorectal (COAD) tumors in the TCGA data set (two-tailed Wilcoxon rank-sum test, p = 2.1e–13). **F)** Gene set dysregulation associated with Chr1q gain in breast single cells: Heatmap showing significantly enriched 'Hallmark' gene sets in Chr1q-gain vs. Chr1q-WT breast tumor cells. Colors represent pooled ssGSEA effect sizes. **G)** Notch signaling is upregulated in Chr1q-gain breast cells: Violin plot comparing 'Hallmark Notch Signaling' gene set scores (via ssGSEA) between Chr1q-gain and Chr1q-WT cells across 7 breast tumors. A significant upregulation is observed (paired two-tailed Wilcoxon signed-rank test, p = 0.047). **H)** Notch2 signaling is significantly upregulated in Chr1q-gain breast cells: Violin plot comparing 'Reactome Signaling by Notch2' gene set scores (via ssGSEA) between Chr1q-gain and Chr1q-WT cells across the same 7 breast tumors. A significant upregulation is observed (paired two-tailed Wilcoxon signed-rank test, p = 0.016). **I)** Upregulation of 'Hallmark Notch Signaling' in TCGA breast tumors with Chr1q gain: GSEA plot showing significant upregulation of the 'Hallmark Notch Signaling' gene set in Chr1q-gain vs. Chr1q-WT breast (BRCA) tumors in the TCGA data set. **J)** Upregulation of 'Reactome Notch2 signaling' in TCGA breast tumors with Chr1q gain: GSEA plot showing significant upregulation of the 'Reactome Signaling by Notch2 pathway*'* in Chr1q-gain vs. Chr1q-WT breast (BRCA) tumors in the TCGA data set.

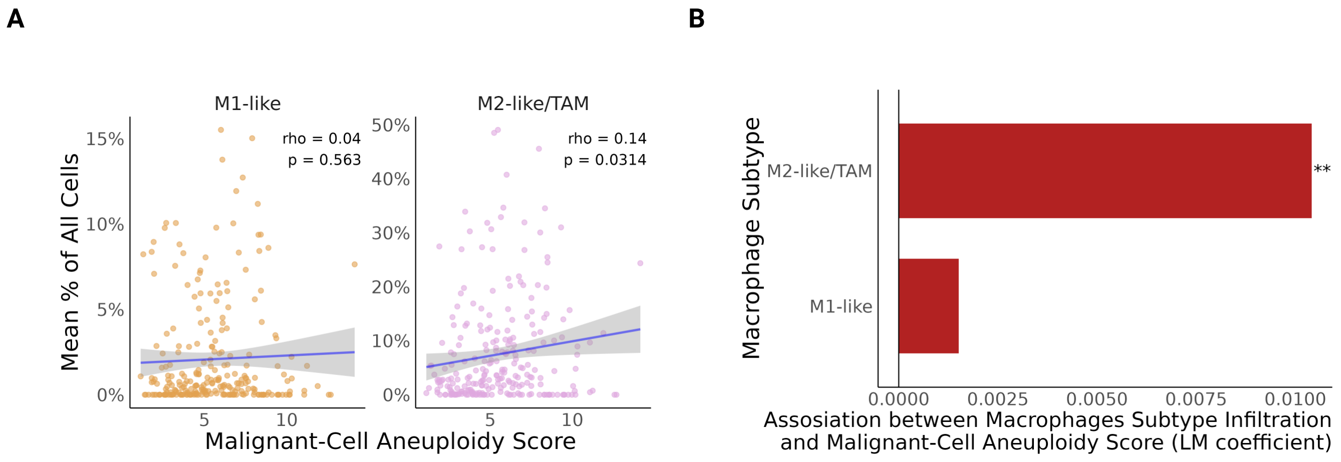
A
M1-like
M2-like/TAM
rho = 0.04
p = 0.563
rho = 0.14
p = 0.0314
15%
10%
5%
0%
50%
40%
30%
20%
10%
0%
5
10
5
10
Mean % of All Cells
Malignant-Cell Aneuploidy Score
B
Macrophage Subtype
M2-like/TAM
M1-like
**
0.0000
0.0025
0.0050
0.0075
0.0100
Assosiation between Macrophages Subtype Infiltration
and Malignant-Cell Aneuploidy Score (LM coefficient)

**Supplementary Figure S15: M2-like/TAM enrichment is consistently associated with malignant-cell aneuploidy (related to Fig.5)**

**A)** Correlation between aneuploidy scores of the malignant cells and macrophage abundance in the tumor: Scatter plots visualizing the relationship between the malignant-cell Aneuploidy Score (AS, x-axis) and the proportion of specific macrophage subtypes (y-axis). Each point represents a tumor (n = 248). A significant positive correlation is observed for the M2-like\TAM macrophages (Spearman's rho = 0.14, p = 0.0314. For M1-like rho = 0.04, p = 0.563). **B)** Association of macrophage subtypes with aneuploidy: Bar plot displaying effect estimates (regression coefficients, x-axis) for each macrophage subtype (y-axis) as a function of the malignant-cell AS. Coefficients were derived from linear regression models controlling for cancer-type-specific confounding effects. Red bars indicate positive (enrichment) associations, with asterisks (*) denoting statistical significance (* q < 0.05, ** q < 0.01, *** q < 0.001).
